# Sensing Echoes: Temporal Misalignment in Auditory Brainstem Responses as the Earliest Marker of Neurodevelopmental Derailment

**DOI:** 10.1101/2022.01.27.478048

**Authors:** Elizabeth B Torres, Hannah Varkey, Joe Vero, Eric London, Ha Phan, Phyllis Kittler, Anne Gordon, Rafael E. Delgado, Christine F. Delgado, Elizabeth A. Simpson

## Abstract

Neurodevelopmental disorders are on the rise worldwide, with diagnoses that detect derailment from typical milestones by 3-4.5 years of age. By then, the circuitry in the brain has already reached some level of maturation that inevitably takes neurodevelopment through a different course. There is a critical need then to develop analytical methods that detect problems much earlier and identify targets for treatment. We integrate data from multiple sources, including neonatal auditory brainstem responses (ABR), clinical criteria detecting autism years later in those neonates, and similar ABR information for young infants and children who also received a diagnosis of autism spectrum disorders, to produce the earliest known digital screening biomarker to flag neurodevelopmental derailment in neonates. This work also defines concrete targets for treatment and offers a new statistical approach to aid in guiding a personalized course of maturation in line with the highly nonlinear, accelerated neurodevelopmental rates of change in early infancy.

**Significance Statement:** Autism is currently detected on average after 4.5 years of age, based on differences in social interactions. Yet basic building blocks that develop to scaffold social interactions are present at birth and quantifiable at clinics. Auditory Brainstem Response tests, routinely given to neonates, infants, and young children, contain information about delays in signal transmission important for sensory integration. Although currently discarded as gross data under traditional statistical approaches, new analytics reveal unambiguous differences in ABR signals’ fluctuations between typically developing neonates and those who received an autism diagnosis. With very little effort and cost, these new analytics could be added to the clinical routine testing of neonates to create a universal screening tool for neurodevelopmental derailment and prodrome of autism.

## Introduction

There is substantial evidence that brain related developmental disorders are present perinatally or earlier (1, 2). However, currently, except for some genetically related disorders, diagnosis does not occur until there is a delay in acquiring developmental milestones, by which time the brain circuitry is largely formed. This precludes “*very early intervention*” which could ameliorate at least some of the pathological course of the disorders (3).

Developmental trajectories traditionally trace physical growth such as weight, height, and head circumference, according to standard charts set by the Centers for Disease Control and Prevention and the World Health Organization (4). Other parameters related to the ways in which infants move, play, emotionally self-regulate, and socially interact and communicate with others, have been adopted as possible observable descriptors to flag departure from expected timely milestones (5, 6). These qualitative criteria, however, miss the opportunity to detect the maturation differences much earlier, as many observable social-communication skills do not emerge until after the first year of life. Their proper scaffolding depends on the timely maturation of building blocks of social interactions, including sensory processing, integration, and the dynamics of sensory-motor transformations. Furthermore, some aspects of detectable behaviors start as normal, becoming abnormal only when they persist into later infancy or beyond (7, 8).

On average, diagnosis of neurodevelopmental disorders coexisting within the spectrum of autism occurs at 4 years and 3 months with 50% of children not diagnosed until age 6. Despite much effort, age at diagnosis has improved little over the past couple of decades (9). Given that earlier therapeutic interventions are associated with better long-term outcomes (10), a challenge that we face is detection of affected children as early as possible and identification of targets for treatment at a very early neurodevelopmental age (7).

Recent biological evidence supports the notion that at least some forms (and probably the majority) of the broad, heterogeneous developmental disorders within the autism spectrum (ASD) are atypical prior to birth, many linked to genetic origins (11–13). These in turn, may have significant effects on postnatal development (14–17). The obstacle to identifying the prenatal changes is a lack of dynamic developmental biomarkers because most of the evidence for prenatal pathology comes from mostly static postmortem cellular, molecular, and genetic findings (2, 18).

An accessible type of assessment that is routinely done on newborns and that could provide dynamic information changing at a micro-level, beyond the observational limits of the naked eye, is the Auditory Brainstem Response (ABR). This test is currently done as a hearing screening test. However, with little extra time or cost needed, we could adapt it into a valid screening tool to forecast neurodevelopmental disorders. A barrier here is the type of methodology currently in use to analyze ABR waveforms and their inherent variability.

The ABR waveform produces a Dirac-delta like peak in response to the input signal consisting of clicks or bursts sound stimuli, and stimulation decibel (dB) level (Figure 1A inset.) Upon repetitions of the sound stimulus (Figure 1A), the waveforms reflecting the ABR are averaged under the theoretical assumption that the variations in parameters of interest (*e.g*., peaks’ amplitudes and latencies) distribute normally, and that as such, their stochastic, moment-to-moment micro-fluctuations are superfluous, beyond one or two (assumed normal) standard deviations of the (assumed normal) mean (Figure 1B). As it turns out, at a micro-level of inquiry, not all such parameters’ fluctuations distribute normally, nor are they stationary. To avoid ignoring such variations, under an *a priori* imposed theoretical mean, and to capture the non-stationary and nonlinear nature of the neonatal data, new analyses call instead for empirical estimation of the most adequate continuous family of probability distributions that can more generally characterize the stochastic properties of the parameters routinely extracted from ABR signals and from other features of the data that have been thus far underutilized (*e.g*., Figure 1C-D).

**Figure 1.**
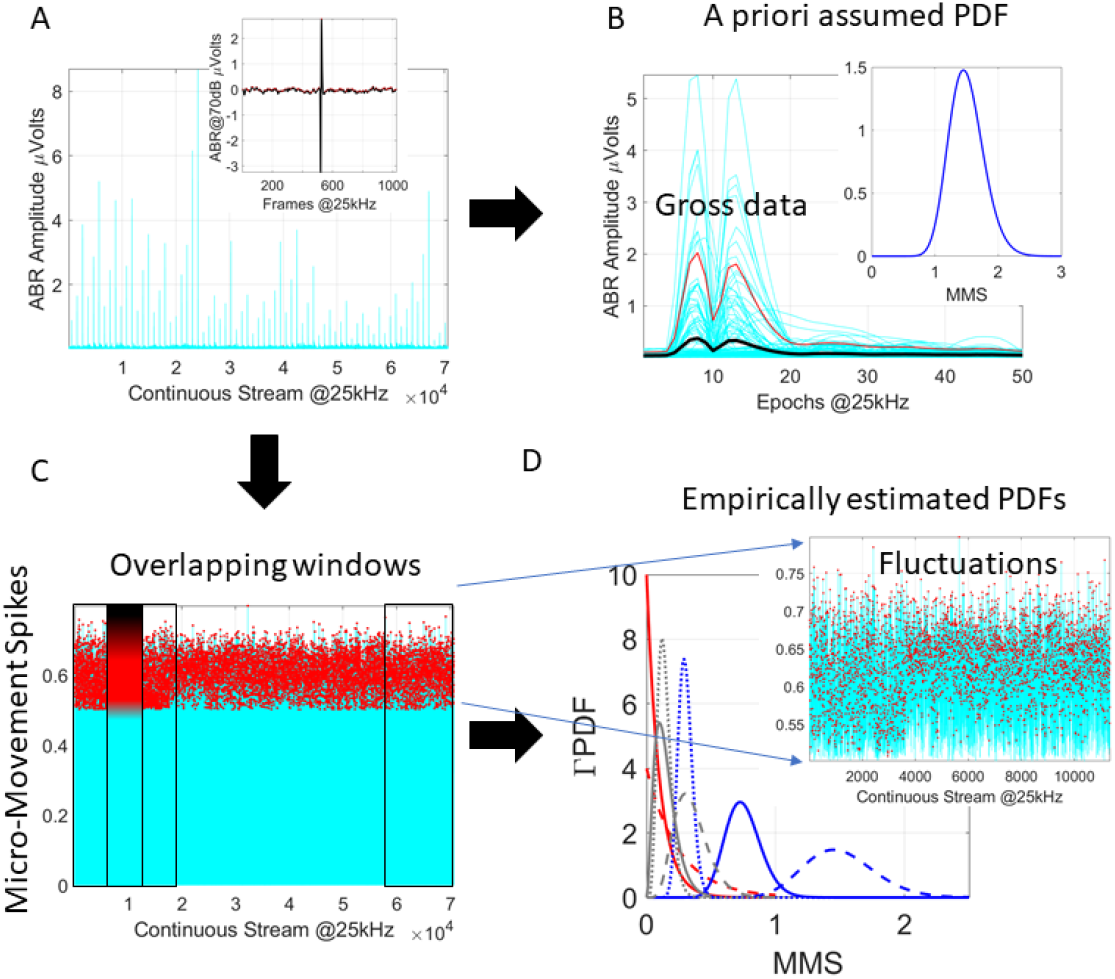
The need for new analytics. (A) Sample data from ABR pooled over multiple trials. Inset shows one trial while time series show the absolute (positively rectified) responses across trials. (B) Traditional approach is to take epochs of the data and stack up the waveforms to obtain the mean and standard deviation under an *a priori* theoretically assumed distribution (*e.g*., the Gaussian distribution in the inset). The theoretical mean is the black curve, and the red curve is two standard deviations from the theoretical mean. The data above the red curve is considered superfluous (gross data) and as such does not enter the analyses. (C) A different approach considering the moment-by-moment fluctuations away from an empirically estimated mean (standardized between 0-1 to scale out disparate anatomical sizes of e.g., head circumference). These fluctuations change from window to window (D), shifting in a nonstationary way according to shifts in probability distribution function (PDF). These PDFs best fit the data in a maximum likelihood estimation (MLE) sense with 95% confidence. Inset zooms in a sample window of fluctuations in C, requiring a minimum of 100 peaks to ensure high confidence in the estimation.

Adopting new methods of analyses that rely on individual empirical fluctuations rather than on theoretically assumed population means would also enable *personalized* characterizations and cross-sectional analyses of changes over time. In this sense, we could standardize the waveforms, to scale out possible allometric effects (*e.g*., differences in head circumference) that emerge in early infancy, from non-linear, rapid, and highly dynamic changes in patterns of physical growth and individual neuroanatomical differences across the population (15, 18), to cross-sectionally examine large cohorts of participants over time.

Evidence from ABR studies suggests that prenatal maturation in the central auditory system (19, 20) far precedes other aspects of sensory-motor maturation, which in turn form the foundation of social interactions and communication. Indeed, as altricial mammals, human babies require a long time of maturation of sensory-motor control (21, 22), yet at birth the brainstem already functions to mediate a few survival activities in the neonate (*e.g*., breathing, swallowing, excreting and crying, perhaps as a first rudimentary form of communicating states of hunger, discomfort, sleepiness, *etc*.)

We combine ABR with clinical data obtained from the neonates and young infants to study anew empirically estimated signatures of micro-variations informed by ASD diagnosis. Under this new personalized statistical framework, we reveal important individual features that automatically and unambiguously forecast departure from typical neurodevelopment at the earliest time upon birth. This digital approach can be easily incorporated into current ABR-based hearing screening tools, to flag neurodevelopmental derailment at birth, and at large scale.

## Methods

### Experimental model and subjects’ details

The research protocol was approved by the Institutional Review Boards of involved institutions and written informed consents were obtained from parents/guardians of all participants.

Three data sets were used in this study spanning from neonatal stages to early childhood. The first two sets are from neonates. The third set spans from 1.8-6.8 years of age. The first set comprises 233,915 neonates from the neonatal intensive care unit (NICU) and the well-baby nursery (WBN) including well balanced numbers of males and females. This set comprises the ABR waveform with clear Peak V latency data and response (Dirac-delta response) waveform as that shown in Supplementary Figure 1B. Only babies with high signal to noise ratio of the output were included with clear peak prominence of the waveform and Peak V latency information available.

In set 2 we had these responses as in Supplementary Figure 1C including three dB levels (70, 75, and 80dB). We divided set 2 into samples denoted L1, babies with multiple trials per dB level, and L2-L3, babies with trials for one or two of the dB levels to be used in pooled data and bootstrapping analyses. For our inclusion criteria, we ensured similar instrumentation and inspected the data for high-quality signals and as variable demographics as possible. Exclusion criteria were noisy and/or incomplete sets of data.

This second set (sample of representative demographics shown in Supplementary Figure 1 and Supplementary Table 1) comprises 54 babies, 30 having received the ASD diagnosis years later. The data available comprised full waveforms for 3dB levels (μV) *vs*. time (recorded at 25kHz) as in Supplementary Figure 2C. The data also had latency peak data for peaks I-VII (*ms*). These are shown in Supplementary Figure 6, where we report individually for each peak, the response latency histograms of ASD *vs*. non-ASD babies. The pairwise *p-values* from statistical comparison are in Supplementary Figure 6H. Supplementary Figure 7 provides the lower triangular similarity matrix (because it is a symmetric matrix) of pairwise comparison of these frequency histograms. These differences are measured using a proper distance metric, the Wasserstein-Kantorovich distance metric (*a.k.a*. the Earth Movers’ Distance metric (23–25) to assess if large effects are present in the data. This set had additional information on birth weight (BW) and estimated gestational age (EGA), which were used in Supplementary Figure 1 to demonstrate the nonlinear (exponential) nature of neonatal growth and motivate the need to standardize the waveform using Equation 1 below. These quantities were also used in other analyses depicted in Supplementary Figures 5A, 8A, 8B and 15. The mean BW in babies who went on to develop typically was 2.1919 × 10^3^ grams (+/- std 1.0739), median 1.9845 × 10^3^ grams *vs*. those who received the ASD diagnosis 1.82 73 × 10^3^ grams (+/- std 988.5), median 1,715 grams. The mean EGA for noASD babies was 34 weeks (+/- std 5.20), whereas the ASD babies had mean EGA 34.5 weeks (+/- std 4.9), median 31.5.

The third set comprised 69 infants with 18 having received the ASD diagnosis and, similarly to the second set, it has a skewness towards males owing to the current deficient pipeline from diagnosis to research (26). In this set we do not have information on whether the babies were pre-term or full-term.

**Methods Table 1.**
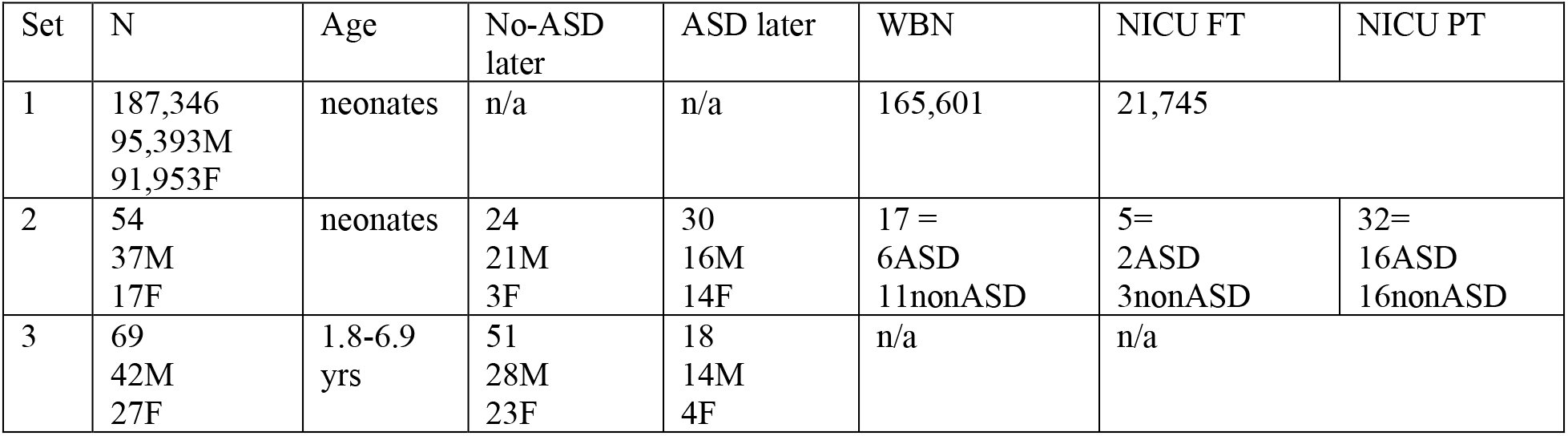
lists the three sets with the demographics and breakdown for the babies by sex (XX-F vs. XY-M) with full set of clean data. In set 1, we include records with full peak prominence data (μV) and Peak V latency. Information on WBN *vs*. NICU was available but no breakdown of PT *vs*. FT was available. In set 2 we include babies with at least 3 full trials inclusive of 70, 75, 80 dB levels of input each. (Additional babies who had 1 or 2 trials for one- or two-dB levels are included in the Supplementary Material Table 1. These were used in pooled-data analyses and in the bootstrapping analyses.) Set 3 includes clean trials only and breaks down data by sex but no information for these infants and young children was available on WBN *vs*. NICU status at birth.

## Quantification and New Statistical Analyses

### Micromovement Analyses

Supplementary Figure 2 explains the derivation of the micro-movement spikes MMS, a general datatype that standardizes the time series waveforms for further analyses that involves diverse populations, while preserving the original information on the time stamps of the peaks and their original amplitude ranges. The notion of micro-movements has been patented by the US and EU Patent offices based on over 20 peer-reviewed papers (not cited here to minimize self-citation) (27). These methods and data are also openly accessible to the scientific community in zenodo, https://zenodo.org/record/6299560#.Y3las3bMLcs.

The MMS turn continuous analog signals to digital spike representations in [0,1] real numbers range. This data type is built from normalized deviations of the original waveform from the empirically estimated mean amplitude. Then these time series of absolute deviations from the mean is normalized using equation 1:

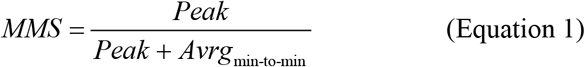

This normalization scales out allometric effects from disparate head circumferences across babies and permit analyses of the noise-to-signal-ratios (NSR) of the original time series data and of their information content. Peak is the local peak value of the waveform, sandwiched between two local minima, then the averaged value of all points comprising the Peak from minima to minima is used in the denominator. This is a real value between [0,1] that nevertheless maintains the original timing of the peak and can be scaled back to its original physical units. Because the information comes from data that is traditionally smoothed out as noise (gross data) in current statistical practices (e.g., Figure 1), MMS analysis tends to non-obviously reveal hidden information that has been missed by well-studied phenomena. The MMS series consists of quiet times of mean activity interspersed with bouts of activity away from the baby’s mean baseline. The signature is personalized. Here we combine the resting state jitter preceding the Dirac-delta like pulse and the jitter from the post-pulse refractory period with the pulse, to then extract the MMS from the full sequence of ABR points collected at 25kHz.

### Stochastic Analyses

Supplementary Figure 3 explains the estimation procedure and shows the analytical pipeline to derive interpretable parameter spaces and to make empirically informed statistical inferences from the digital MMS data. This method enables optimal feature identification and parameter spaces to stratify phenomena in general and uncover important patterns in the data offering potential targets for treatments. Similarity metric (Earth Mover’s Distance, EMD (24)) can be used to measure distances between points in abstract probability space and investigate the rates of stochastic shift. in Supplementary Figure 3A frequency histograms can thus be easily compared.

## Results

In the first cohort of neonates (187,346 viable data / 233,915 records), we found large differences in the empirically estimated probability distribution functions (PDFs) of the peak V latency between WBN and NICU babies (two-sample Kolmogorov-Smirnov test, p<0.01). Cross-sectionally, WBN neonates showed systematic trends in decreasing peak latency over 8 weeks post birth. In stark contrast, NICU neonates’ peak latencies remained stagnant throughout this time. This is shown in Figure 2A.

**Figure 2.**
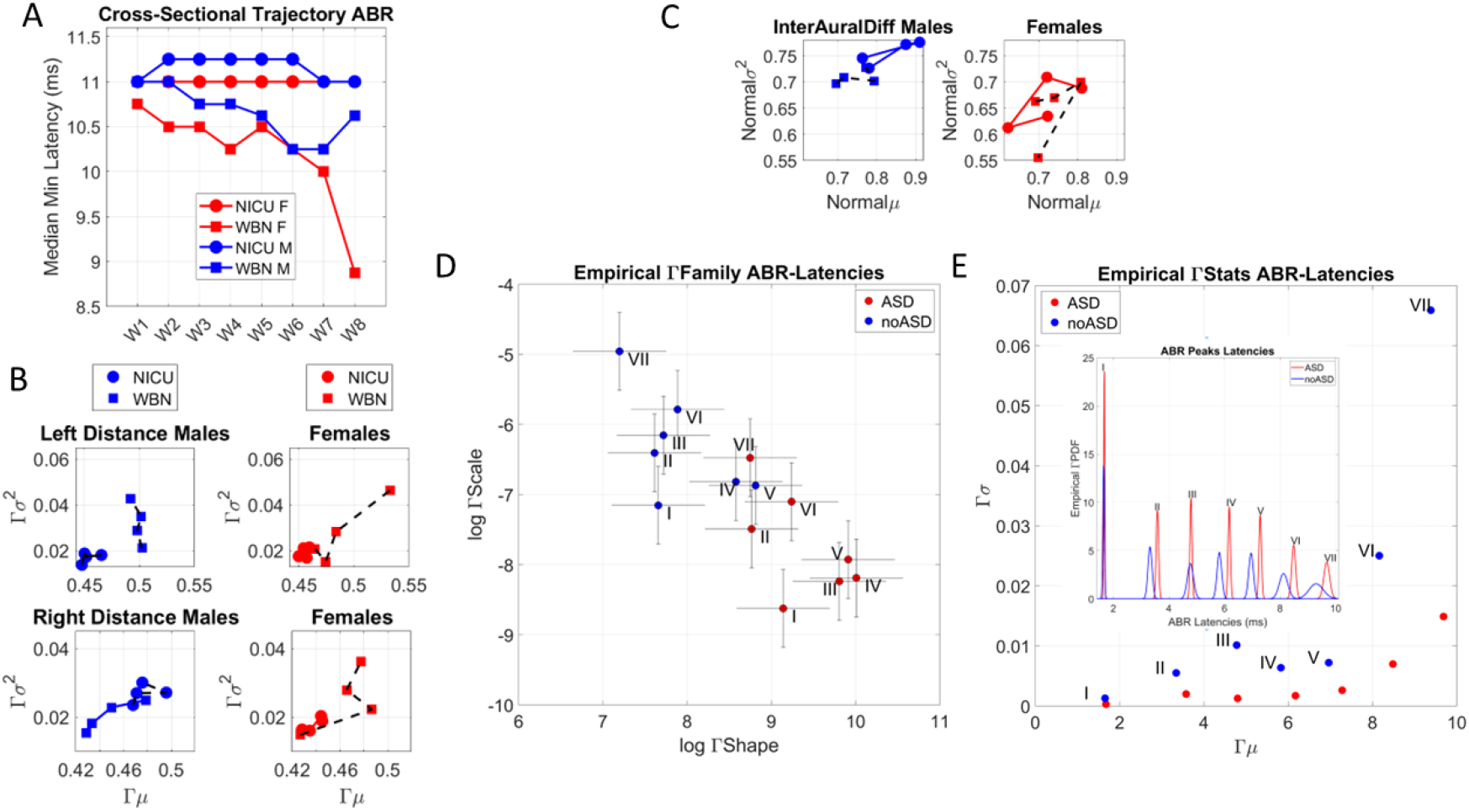
Differentiating neonates’ neurodevelopment and those resulting in an autism diagnosis. (A) Neonatal Intensive Care Unit (NICU) *vs*. Well Baby Nursery (WBN) neonates received the ABR test. Cross-sectional data reflecting the median minimum latency (in milliseconds) of peak V, taken across babies who received the test at each of 8 weeks after birth in the set of 230+K neonates. NICU babies tend to maintain the same level of latency, while WBN babies are trending downward, towards faster responses as weeks go by, also distinguishing between males and females. (B) Examination of patterns of variation in the waveforms’ amplitude from the right and left ear, for 100 randomly selected neonates in each of the first 4 weeks, separates the cross-sectional trajectories of NICU *vs*. WBN and distinguish between males and females. Distance refers to the length of the waveform, taken as the cumulative distance from adding each delta value across time of the test. Deltas are obtained from differentiating the smooth curve representing the ABR waveform. These results extended to the cross-sectional evolution of peak V latencies and their absolute right-left difference denoting interaural time differences (C). (D) Delayed response latencies and narrow bandwidth of latency ranges in neonates that later receive the ASD diagnosis. The second neonatal data set, with babies that received an ASD diagnosis, show families of empirically estimated continuous Gamma probability distribution functions (PDFs). These emerge from the latencies of the response potential and differ for the ASD *vs*. non-ASD neonates across each of the I-VII regions. Log-log Gamma parameter plane spanned by the Gamma shape and scale parameters show for each brainstem region a complete separation between cohorts. Points represent the PDF’s shape and scale (dispersion), with 95% confidence intervals. The ASD-neonates localize in the region of distributions with lower dispersion and more symmetric shapes. (E) The empirically estimated Gamma moments (mean and variance) emphasize the differences in (C). The non-ASD neonates have a broader range of latency variability and faster timings than the ASD-neonates. Inset shows empirical PDFs superimposed for both cohorts, highlighting the shifts in timings (ms) as the response signals propagate along the seven regions.

This cross-sectional trajectory suggests NICU infants may have delays in decreasing latencies, signaling a lack of improvement in transmission speed found in healthy babies over the 8 postnatal weeks. The results were consistent for male and female neonates, but WBN females showed a faster trend in latency reduction, toward faster response propagation times (*ms*). However, this differentiation was far more modest in the female NICU babies. Supplementary Figure 4 shows their stochastic shifts across the first 4 weeks of life.

Further analyses of the waveform were carried out by differentiating the response time series comprising 61 and 62 points (0.2-ms period) for the left and right ear signals, respectively, and taking the cumulative sum of these values. This information revealed the length of the curve that we obtain from the differentiation. The parameter indicates patterns of amplitude variability for each group. Shorter lengths spanned by taking this cumulative sum of the delta values from the differentiation of the original smooth ABR waveform (the ABR curve) indicate lower variability and lower amplitude. We thus obtained all the waveform lengths per group and randomly selected 100 neonates in each group, to empirically estimate the family of distributions best characterizing the patterns of waveform length variability, in an MLE sense. We coin the waveform length “Distance” and obtain it for the left and right ears and for males and females (Figure 2B). The continuous Gamma family of probability distributions was the best fit for this parameter. It revealed fundamental differences in such patterns, depicted in Figure 2B along a parameter plane spanned by the empirically estimated Gamma moments (mean and variance). These are shown for the right and the left ear and for the males and females of each of the ASD and non-ASD groups and for each of the first 4 weeks of life (taken cross-sectionally). We see the shorter trajectories of the NICU babies, denoting a profound lack of variability in the waveform and lower amplitude signal than the WBN neonates (Wilcoxon rank sum test, p<0.01).

We then obtained, for each group and week, the interaural differences by subtracting the latencies at which peak V were attained. This analysis revealed marked differences in the cross-sectional trajectories spanned by the mean *vs*. variance values, which were best fit by a normal distribution, according to MLE. These are shown in Figure 2C.

Such detectable differences across WBN and NICU babies during the first weeks of life prompted the analyses of the second data set, which also included Full Term (FT) and Pre-Term (PT) babies in the NICU and in the WBN. We asked if fundamental differences in delayed latencies could be found in this new cohort of babies who were clinically tracked for the next four years.

Importantly, among these babies, we had a group with neurodevelopmental disorders that went on to receive a diagnosis of ASD later, after 4 years of life. These ASD babies were compared to a group that developed along a typical path and did not receive the diagnosis, (non-ASD.) The peaks’ latency data were examined for all I-VII peaks using empirical estimation and distributional analyses of peaks’ fluctuations in amplitudes and latencies, as explained in Supplementary Material Figure 3. More precisely, for each parameter of interest (peaks’ latencies (ms) and waveforms’ features (peaks’ amplitudes and prominences (μV) and peaks’ widths (ms)), we derived the micro-movements spikes MMS standardized waveform, which scales out allometric effects due to disparities in head circumference across babies. This normalization of the waveforms is important to ensure appropriate comparisons across different ages and rates of growth during early neonatal neurodevelopmental stages. To demonstrate the non-linear rate of growth through the body weight (g), we show in Supplementary Figure 1A representative babies of FT and PT males and females with ASD and non-ASD types, while panel B shows the exponential fit taken cross-sectionally over the span of 24-41 weeks of estimated gestational age (EGA) for both ASD and non-ASD sets.

Figure 2D shows the results of our empirical fit to the I-VII peaks’ latencies expressed on the Gamma parameter plane. The continuous Gamma family of probability distributions was the best fit to the peaks’ latency data, in an MLE sense. As such, each point on the Gamma parameter plane represents the empirically estimated stochastic signatures corresponding to the latency values of each of the ASD and non-ASD groups when each peak (I-VII) was detected. The ASD (red) and non-ASD (blue) span different PDFs for each peak. These PDFs are plotted as points on the Gamma parameter plane spanned by the shape axis and the scale axis. Each point is an estimated PDF with the 95% confidence intervals for each of the shape and scale (dispersion) parameters of the continuous Gamma family of probability distributions. This family was the best fit in an MLE sense (over other distributions such as the lognormal, the normal, the Weibull and the exponential), for the frequency histograms of the latency corresponding to each peak and group. We note that there are unambiguous differences between the ASD and non-ASD groups for each of the 7 peaks. Furthermore, we note that in Figure 2E where we plot the PDFs along the continuous timeline (ms) the ASD group is systematically lagging, with cumulative delays reflecting how the response to the pulse propagates along the brainstem sites.

We also highlight the difference in distribution dispersion (width) signaling broader ranges of latencies in non-ASD group and a pronounced reduction in latency range in the ASD group. This implies a much narrower bandwidth in processing sounds’ frequencies for the ASD group. The richer variability and systematically shorter latencies of the non-ASD group can also be appreciated in Figure 2E, where we plot in the inset, the plane spanned by the empirically estimated first two Gamma moments (mean and variance) corresponding to the PDFs in the inset. To better see these results, we plot at scale each empirically estimated Gamma PDF for each peak along the anatomical regions of the brainstem.

Figure 3 displays each PDF per peak, to show the differences in propagation delays more clearly. The cumulative effect across these sites of the brainstem revealed 1.74*ms* delay, with 0.40*ms* as the longest local delay for ASD in the region VII, while region III had the shortest delay 0.05*ms*. Supplementary Table 2 summarizes all latencies (*ms*) across the seven points of interest for each participant type. The ASD neonates show a net cumulative latency of 8*ms vs*. 7.62*ms* in the non-ASD neonates. Considering that sound processing occurs at microseconds time scale, these delays in latencies found in the ASD babies are very large. We later discuss potential ramifications of these persistent delays and narrow bandwidth of latencies range found across all regions of the brainstem of the ASD babies.

**Figure 3.**
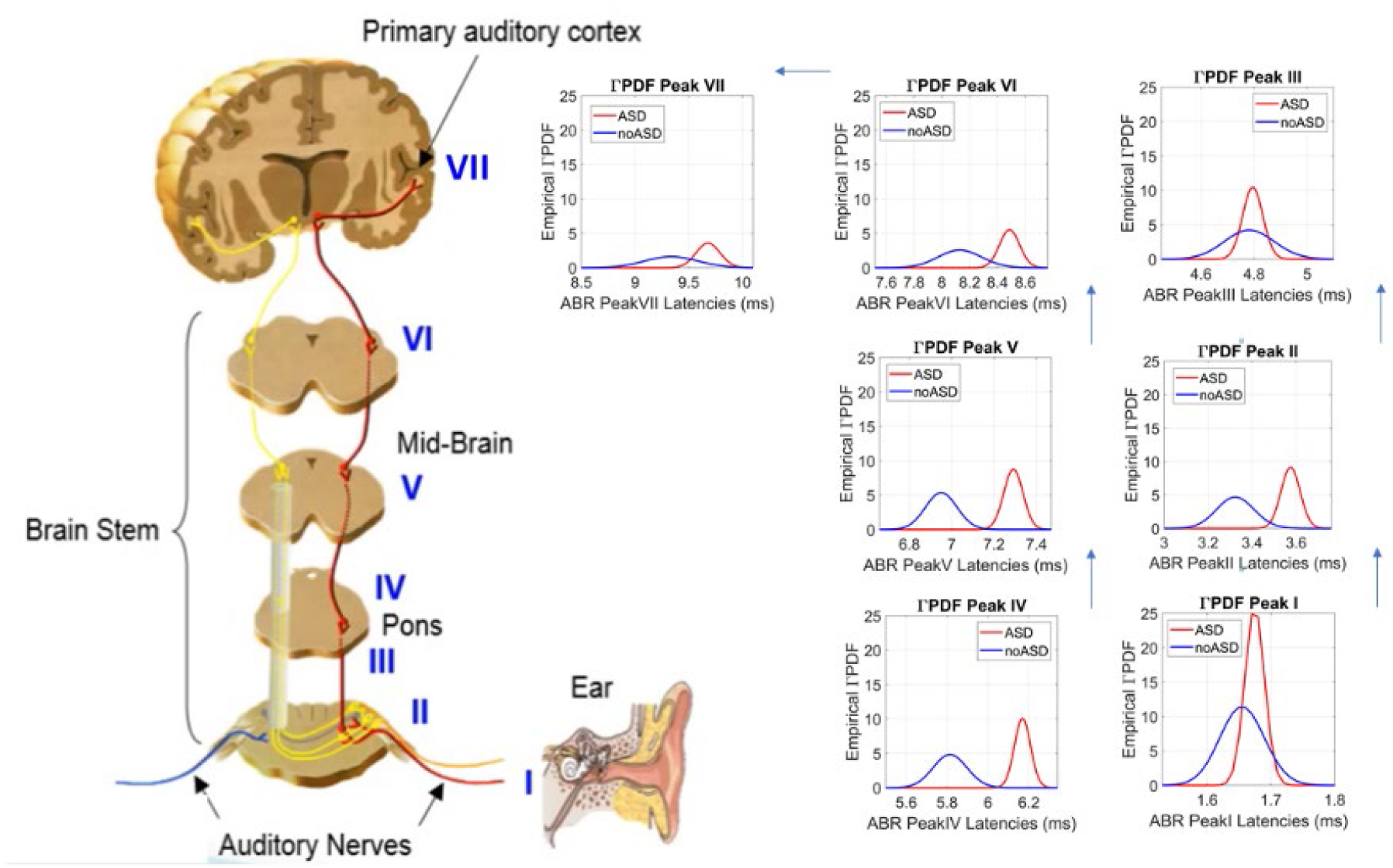
Zooming into the empirically estimated family of continuous Gamma probability distributions characterizing the ASD and non-ASD cohorts of 54 neonates. Each plot compares the two ASD *vs*. non-ASD PDFs for a site (I-VII) and clearly demonstrates the (cumulative) delays in the ASD group along with the narrower bandwidth of values. Brainstem schematic is included for visualization purposes.

These results are expanded in Supplementary Figures 5-7, where we perform non-parametric statistical comparisons across group types. For example, Supplementary Figure 5A expands the systematic shifts in peak latencies for males and females in each of the ASD and non-ASD groups while also providing information on the body weight (g) proportional to the marker size. This is also done for the Bayley’s score ranges in Supplementary Figure 5B. Importantly, Supplementary Figure 6A-G provides frequency histograms of latencies for each peak and ASD vs. non-ASD. The comparison matrix in panel H, is derived using non-parametric Kolmogorov-Smirnov test via bootstrapping by drawing from the larger set the number of measurements of the smaller set and forming 100 distributions. The median *p-value* is obtained to construct a pairwise matrix of comparisons for each of the 7 sites and for the ASD vs non-ASD groups. This matrix is then color-coded by scalar p-value and entries with two asterisks are significant p <0.01, while those with one asterisk are p<0.05. Supplementary Figure 7 then expresses the lower-triangular matrix comparing pairwise the frequency histogram of such latencies in I-VII peaks representing the brainstem regions. To that end, we employ a proper distance metric to measure the differences between frequency histograms and color code the entries of the matrix accordingly (using the normalized value of the Earth Movers’ Distance.)

Supplementary Figure 8 expands from cross-sectional population to a personalized approach by localizing individual’s body weight (g) and sex signatures for each group. These plots are also informed by clinical scores and waveforms’ features that were never examined before. These include micro-fluctuations before and after the Dirac-delta (explained in Supplementary Figure 2) and converted to MMS time series of normalized peaks’ amplitudes, prominences, and widths, whereby we scale out allometric effects of e.g., disparity in head circumference. We note here that these results are skewed toward ASD males due to the insufficient number of females in any random draw of the population (current ratio of 4.5-5 ASD males per each ASD female (28)).

Further analyses of the waveform’s features (peaks’ prominences, amplitudes, and widths also separated ASD from non-ASD in full-term *vs*. pre-term babies. Supplementary Materials Figure 9 shows the unambiguous distinction of dB levels (important to assess responses with precision for therapeutic design.) Supplementary Figures 10-12 show that these results extend to other features of the peaks. Besides prominences (identified as the best feature to separate subgroups), the amplitude and widths also serve that purpose. Results from non-parametric comparisons across dB levels in Supplementary Material figure 13 show significant differences according to the Wilcoxon rank sum test.

### ABR Forecasting ASD in Early Infancy and Childhood Development

As revealed by the Supplementary Materials Figure 1, during early neurodevelopment of the neonate, there is a highly non-linear relation between BW and EGA. As such, assessing the ABR parameters as the system grows and matures past the first year of life, seemed important (21, 22). It is possible that with the maturation and growth of the nervous system (e.g., increased myelination patterns) such disparities in delayed latencies between ASD and non-ASD children disappear. Alternatively, we may still find differences during early infancy and early childhood.

To test these propositions, we had access to the third cohort of ABR data like that of the neonates but involving instead young infants and young children between 1.8 and 6.8 years of age. Importantly, in this group, 18/65 children received a diagnosis of ASD, thus providing referencing criteria for the detection of neurodevelopmental derailment. We had access to multiple trials per child with high signal-to-noise ratio and clear waveform pattern collected with 40kHz sampling resolution. Demographic information (female *vs*. male) was also available, thus facilitating comparisons. Unambiguous statistically significant differences were found, consistent with those detected in the neonates. The empirically estimated stochastic parameters of the continuous Gamma family of distributions are depicted in Figure 4 using a similar format as before.

**Figure 4.**
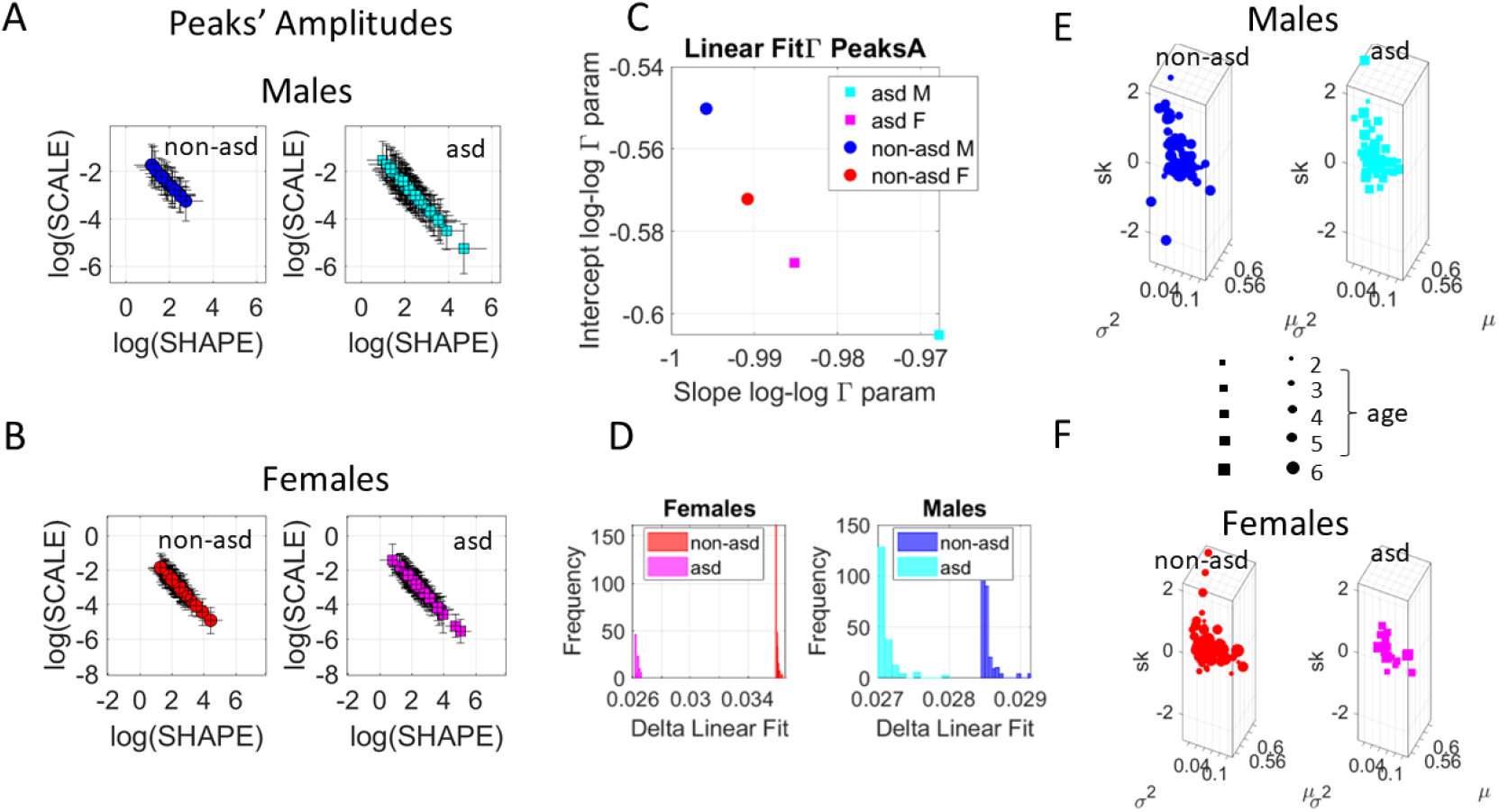
Differentiating the peaks’ prominences in the ABR from typically developing (TD) young children (1.8-6.8 years old) and those with an ASD diagnosis. (A) Gamma PDF parameters empirically estimated using micro-movement spikes derived from fluctuations in the waveform in males vs. (B) females. (C) Slope vs. intercept fitted to the scatters in A-B differentiate males from females within and between the groups, and ASD from non-ASD children. The histograms of the delta value capturing the goodness of the linear fitting to the log-log scatter, separate each ASD from non-ASD across the groups of females and males (D-E) Corresponding Gamma moments (mean, variance, and skewness) empirically obtained from the Gamma shape and scale MLE parameters in A-B and sized according to age in legend, permit visualizing the individualized stochastic signatures.

Figures 4A-D show the results from our stochastic analyses of the waveforms’ features. Peaks’ amplitude derived from differentiating the original waveform and converting it to the standardized MMS, had microfluctuations revealing marked differences between the ASD and non-ASD groups. ASD had a broader variance over mean ratio, along the scale axis. This differentiation was accentuated by the narrower bandwidth of Gamma distribution shapes for non-ASD (along the shape axis) that extended to males vs. females within and between the groups.

To further explore subgroup differences, we found a linear relationship between the log shape and log scale Gamma parameters (Figure 4A-B). These spanned over 2 decades of the log horizontal axis, suggesting a power law like relation. This relation prompted us to compute the slope and intercepts of the line fitting the log-log scatter. A parameter plane spanned by these two scalar quantities served to localize each cohort, thus revealing unambiguous differences between ASD and non-ASD in general, but also differences between males and females in each of the cohorts. We note here in Figure 4 that the differences between males’ ASD and non-ASD are much broader than that of the females’ counterparts. The distributions of the delta value measuring the goodness of linear fitting to the log-log scatter in panels 4A-B and comprising all trials from all children, revealed in Figure 4D unambiguous differences for each child’s trials in each group. Other features—peaks’ widths (ms) related to timing and peaks’ amplitudes from the baseline value (μV)—revealed congruent patterns (not shown here for brevity). Across the population of young infants and young children of this study, differences in ABR parameters were systematically evident between ASD and non-ASD. Individual differences were also captured in panels E-F of Figure 4, where we plot the marker size proportional to age. These results suggest that across the population, differences found in neonatal stages remain at early infancy and early childhood stages.

## Discussion

The ABR constitutes one of the most reliable measures of neural integrity in the cochlear nerve and caudal brainstem pathway, here indicating as well cumulative delays as the sound wave propagates and arrives at the primary auditory cortex in latency VII. In this work, by considering anew the full ABR waveform and its micro-fluctuations over repeated trials, we reveal specific information appropriate to infer the neurological and audiological status of healthy and special-risk populations. Using new analyses on the gross data that is currently discarded as superfluous, we found that WBNs show a significant reduction in the latencies of the peak V during the first two months of life. This faster processing of sound is accompanied by smaller interaural time differences and broad ranges of variability and amplitudes of their responses whereby males’ and females’ trends are highly differentiable early in life. In contrast, this maturation path is different in NICU babies. They remain stagnant, deprived of a reduction in latency delays for peak V that is also accompanied by narrower ranges of variability and lower amplitude of the response waveform. Babies that go on to receive an ASD diagnosis show profound delays in the latencies of the ABR across all 7 regions of the brainstem. Furthermore, these neonates showed systematically narrower range of latencies for each peak, suggesting poor access to the full sound frequency spectrum. Thus, the sensory signal transmission across the caudal brainstem is not only delayed in prodrome ASD. It is also impoverished in those infants and young children who years later received the diagnosis. This information obtained at neonatal stages was consistently confirmed (cross-sectionally) at infancy and early childhood stages, suggesting the need for early detection of the problem and early intervention to steer the system toward typical ranges. These new analyses could also separate male and female participants at this early age. This is important, given the male-to-female ratio disparity acknowledged by current diagnostic criteria of ASD (29).

The present results non-trivially expand ABR analyses providing a personalized statistical platform amenable to incorporate into existing hearing screening tools, to perform the same assessment while screening for possible neurodevelopmental derailment, without much extra time or cost. Given that ASD comprises so many comorbid disorders today, and that there is no screener that captures the non-linear dynamical nature of early neurodevelopment, the present approach would make for a valid screening tool to forecast neurodevelopmental disorders in general. Importantly, cross-sectionally, these differences in ABR parameters remained beyond the first year of life. As such, they are consistently detectable throughout critical periods of cognitive development and physical growth, inclusive of entry-level school year.

Important prior work had proposed ABR signals as an early biomarker flagging neurodevelopmental issues (30–34) but no methods for personalized precision phenotyping at this early age had been provided that could also identify specific targets for treatment and track their non-stationary statistical shifts over time. Here, by considering gross data that is traditionally treated as superfluous, we were able to assess fundamental differences in the peaks’ latencies and waveform’s features related to amplitude and width. Indeed, peaks’ prominences proved to be the most informative, although amplitude and width of the peaks also revealed differences across the groups. We found systematic shifts in peaks’ latency suggesting delayed arrival of the sound at each point of interest along the auditory pathway, throughout the brainstem. Such cumulative delays were accompanied by a significantly narrower range of latencies, suggesting narrower access to the bandwidth of frequencies experienced by those neonates and later in life, by the cohort of toddlers and young children.

There are several important implications of these results. One is that they provide specific and new targets for early intervention. Perhaps through sound stimulation, and while aiming at (*i*) broadening the range of latencies and (*ii*) shifting the delays of transmission at each specific peak towards neurotypical ranges, we could help the system with the ASD signature align those delays to neurotypical ranges. This would be important for communication purposes, as with a cumulative average of 1.72 *ms* delay in sound processing, there is no coincidence in transmission and reception of sound signals between two systems (*e.g*., between a neurotypical and an ASD system). Furthermore, at processing timescale of microseconds, millisecond time delays would prevent having an anchor for proper integration of disparate sensory transduction delays, inclusive of vision, proprioception, and inertial motor time delays, among other sources that the baby’s brain needs to predict and compensate for to dynamically interact with others.

Another advantage of having such specific breakdown in ABR parameters for peaks I-VII is that it may be possible using these personalized analytics, to specifically target each of the 7 regions to (*i*) subtype neurodevelopmental disorders accordingly and (*ii*) to broaden and to shift the delays using a multiplicity of inputs. Lastly, the specificity in detection and differentiation of dB levels facilitates designing of training assays for interventions tailored to the individual’s processing speeds at each of the I-VII regions. Delayed sound processing could interfere with the integration of other sensory inputs as well as with sensory-motor transformations necessary to plan and to perform well controlled purposeful actions, to predict their consequences, and to adapt and generalize them to new environments. These are all core building blocks of social interactions and communication, which become evident and detectable by observation only after 3 or 4 years of age, sometimes even after 6 years of age. In this sense, there is no need to wait so long to detect a problem that is present very early in life and has potential mechanistic causal link to higher order social, communication and emotional behaviors. The latter are precisely the current criteria for diagnosis.

The type of delayed sound processing under narrow latencies’ ranges found here could indeed interfere with the development of social interactions and bring uncertainty and anxiety to the system, thus impacting emotional development as well. During social interactions, the person’s biorhythmic activities, including those dependent on sound processing and integration with other sensory inputs, need to be automatically synchronized for proper communication and mutual timely exchange of verbal and non-verbal, gestural cues. Furthermore, the development of a predictive code (35) to anticipate others’ social actions and their consequences, to timely react and smoothly flow during the interaction, could also be impeded owing to such sound processing delays under narrow and low signal information, persistently found here across the population from birth to early childhood. For all these important reasons, it is possible that these early differences in sound processing could constitute targets for very early intervention.

Prior work across the human lifespan had unveiled scaling laws of voluntary (35) and involuntary (36) motor maturation that are violated in ASD. Scaling laws relating human infant’s rates of growth with rates of neuromotor control development were also found in babies who went on to develop neurotypically, violated by three months of age in those who experienced neurodevelopmental derailment (6). Those ontogenetically related scaling laws of human’s nervous systems’ biorhythmic maturation are congruent with phylogenetic scaling laws relating brain weight and time to walk (21). As altricial (rather than precocious) mammals, human babies require the longest time to mature upright walking patterns. Here we reproduce scaling laws relating probability distribution parameters characterizing temporal- and amplitude-related fluctuations of brainstem biorhythmic response signals. We then use these empirically estimated parametric relations to learn about normative ranges and to interrogate their departures in babies who later received the ASD diagnosis. Using these analyses anew, we detected very early differences as well between males and females in each group.

## Conclusion

Keeping in mind the ontogenetically orderly timeline of the human species’ early neurodevelopment will be key to early detection of departures from neurotypical development with direct consequences impacting social interactions and communication. A neonatal screener of neurotypical development is now within our grasp. Any departure from normative ranges is thus unambiguously detectable at birth.

## Author contributions

EBT conceived study, analyzed data, wrote paper, supervised study, HV curated and compiled data, JV curated and compiled data and created repository for public use of data and code, edited paper, EL, HP, PK, AG designed, collected, curated and analyzed clinical infant and ABR data, edited paper, PK and HP supervised study, RED, CFD, and EAS recruited participants, curated and compiled data, provided resources, and edited the paper, All authors agreed to the final version of the paper.

## Acknowledgements

The research data collection was supported with funds from: New York State Office of People with Developmental Disabilities (OPWDD) and New York State Institute for Basic Research in Developmental Disabilities (IBR); NICHD # P01-HD047281 and Autism Speaks # 7598 (Gardner); National Institute on Deafness and Other Communication Disorders (NIDCD) Small Business Innovation Research (SBIR) # 1R43DC018430–01 (to RED); National Science Foundation (NSF) CAREER Award # 1653737 (to EAS). EBT were partly supported by an endowment from the Early Career Development Award to EBT (2014-2017) by the Nancy Lurie Marks Family Foundation and by the New Jersey Governor’s Council for Autism CAUT18ACE015.

## References

1. Bonnet-Brilhault F, et al. (2018) Autism is a prenatal disorder: Evidence from late gestation brain overgrowth. Autism Res 11(12):1635–1642.

2. Courchesne E, Gazestani VH, & Lewis NE (2020) Prenatal Origins of ASD: The When, What, and How of ASD Development. Trends Neurosci 43(5):326–342.

3. Blanc R, et al. (2021) Early Intervention in Severe Autism: Positive Outcome Using Exchange and Development Therapy. Front Pediatr 9:785762.

4. Kuczmarski RJ, et al. (2000) CDC growth charts: United States. Adv Data (314):1–27.

5. Sheldrick RC, Marakovitz S, Garfinkel D, Carter AS, & Perrin EC (2020) Comparative Accuracy of Developmental Screening Questionnaires. JAMA Pediatr 174(4):366–374.

6. Torres EB, Smith B, Mistry S, Brincker M, & Whyatt C (2016) Neonatal Diagnostics: Toward Dynamic Growth Charts of Neuromotor Control. Front Pediatr 4:121.

7. Wolff JJ & Piven J (2021) Predicting Autism in Infancy. J Am Acad Child Adolesc Psychiatry 60(8):958–967.

8. McCarty P & R.E. F (2020) Early detection and diagnosis of autism spectrum disorder: Why is it so difficult? Seminars in Pediatric Neurology: 100831.

9. Maenner MJ SK, Baio J, et al. (2020) Prevalence of Autism Spectrum Disorder Among Children Aged 8 Years — Autism and Developmental Disabilities Monitoring Network, 11 Sites, United States, (Morbidity and Mortality Weekly Report MMWR Surveill Summ 2020), (CDC).

10. Zwaigenbaum L, et al. (2015) Early Intervention for Children With Autism Spectrum Disorder Under 3 Years of Age: Recommendations for Practice and Research. Pediatrics 136 Suppl 1:S60–81.

11. Torres EB (2021) Precision Autism: Genomic Stratification of Disorders Making Up the Broad Spectrum May Demystify Its “Epidemic Rates”. J Pers Med 11(11).

12. Gunter C, et al. (2022) Heritability of social behavioral phenotypes and preliminary associations with autism spectrum disorder risk genes in rhesus macaques: A whole exome sequencing study. Autism Res 15(3):447–463.

13. Escher J, et al. (2021) Beyond Genes: Germline Disruption in the Etiology of Autism Spectrum Disorders. J Autism Dev Disord.

14. Hazlett HC, et al. (2017) Early brain development in infants at high risk for autism spectrum disorder. Nature 542(7641):348–351.

15. Hazlett HC, et al. (2011) Early brain overgrowth in autism associated with an increase in cortical surface area before age 2 years. Arch Gen Psychiatry 68(5):467–476.

16. Hutsler JJ & Casanova MF (2016) Review: Cortical construction in autism spectrum disorder: columns, connectivity and the subplate. Neuropathol Appl Neurobiol 42(2):115–134.

17. Wegiel J, et al. (2013) Contribution of olivofloccular circuitry developmental defects to atypical gaze in autism. Brain Res 1512:106–122.

18. Bauman ML & Kemper TL (2005) Neuroanatomic observations of the brain in autism: a review and future directions. Int J Dev Neurosci 23(2-3):183–187.

19. Ponton CW, Eggermont JJ, Coupland SG, & Winkelaar R (1992) Frequency-specific maturation of the eighth nerve and brain-stem auditory pathway: evidence from derived auditory brain-stem responses (ABRs). J Acoust Soc Am 91(3):1576–1586.

20. Starr A, Amlie RN, Martin WH, & Sanders S (1977) Development of auditory function in newborn infants revealed by auditory brainstem potentials. Pediatrics 60(6):831–839.

21. Garwicz M, Christensson M, & Psouni E (2009) A unifying model for timing of walking onset in humans and other mammals. Proc Natl Acad Sci U S A 106(51):21889–21893.

22. Grillner S & El Manira A (2020) Current Principles of Motor Control, with Special Reference to Vertebrate Locomotion. Physiol Rev 100(1):271–320.

23. Monge G (1781) Memoire sur la theorie des deblais et des remblais. Histoire de l’ Academie Royale des Science; avec les Memoires de Mathematique et de Physique;, (De L’imprimerie Royale, Paris, France).

24. Rubner Y, Tomasi C, & Guibas LJ (1998) Metric for Distributions with Applications to Image Databases. Proceedings of the ICCV.

25. Stolfi J & Guibas LJ (2000) The earth mover’s distance as a metric for image retrieval.. Int. J. Comput. Vis. 40:99–121.

26. D’Mello AM, Frosch IR, Li CE, Cardinaux AL, & Gabrieli JDE (2022) Exclusion of females in autism research: Empirical evidence for a “leaky” recruitment-to-research pipeline. Autism Res 15(10):1929–1940.

27. Torres EB (2018) USPTO.

28. Loomes R, Hull L, & Mandy WPL (2017) What Is the Male-to-Female Ratio in Autism Spectrum Disorder? A Systematic Review and Meta-Analysis. J Am Acad Child Adolesc Psychiatry 56(6):466–474.

29. Mandy W, et al. (2012) Sex differences in autism spectrum disorder: evidence from a large sample of children and adolescents. J Autism Dev Disord 42(7):1304–1313.

30. Baizer JS (2021) Functional and Neuropathological Evidence for a Role of the Brainstem in Autism. Front Integr Neurosci 15:748977.

31. Karmel BZ, Gardner JM, Zappulla RA, Magnano CL, & Brown EG (1988) Brain-stem auditory evoked responses as indicators of early brain insult. Electroencephalogr Clin Neurophysiol 71(6):429–442.

32. Miron O, et al. (2021) Prolonged Auditory Brainstem Response in Universal Hearing Screening of Newborns with Autism Spectrum Disorder. Autism Res 14(1):46–52.

33. Seif A, Shea C, Schmid S, & Stevenson RA (2021) A Systematic Review of Brainstem Contributions to Autism Spectrum Disorder. Front Integr Neurosci 15:760116.

34. Delgado CF, Simpson EA, Zeng G, Delgado RE, & Miron O (2021) Newborn Auditory Brainstem Responses in Children with Developmental Disabilities. J Autism Dev Disord.

35. Torres EB, et al. (2013) Autism: the micro-movement perspective. Front Integr Neurosci 7:32.

36. Torres EB, Caballero C, & Mistry S (2020) Aging with Autism Departs Greatly from Typical Aging. Sensors (Basel) 20(2).

37. Kittler PM, et al. (2011) The development of selective attention and inhibition in NICU graduates during the preschool years. Dev Neuropsychol 36(8):1003–1017.

38. Cohen IL & Sudhalter V (2005) The PDD Behavior Inventory. in Lutz, Fl (Psychological Assessment Resources, Inc., New York).

39. Seethapathy J, Boominathan P, Uppunda AK, & Ninan B (2018) Auditory brainstem response in very preterm, moderately preterm and late preterm infants. Int J Pediatr Otorhinolaryngol 111:119–127.

