## Supplementary materials for "Sensing Echoes: Temporal Misalignment in Auditory Brainstem Responses as the Earliest Marker of Neurodevelopmental Derailment"

### Table of Contents:

- **Information about participants**
- **Supplementary Table 1.** Demographics – number of neonates included in the study with ASD diagnosis
- **Supplementary Figure 1**. Demographic plots for neonates with ASD diagnosis
- **Supplementary Figure 2**. Methods to derive micromovement spikes from ABR signals
- **Supplementary Figure 3.** Methods to study Gamma process
- **Supplementary Table 2.** Results – peak latencies for each site in the two cohorts and inter-peak latency differences.
- **Supplementary Figure 4**. Tracking neonates from NICU and WBN cross-sectionally 4 weeks reveals differentiation between males and females withing and between each group.
- **Supplementary Figure 5**. Empirical Gamma means vs. Peaks' latencies by body weight and Bayley's scores
- **Supplementary Figure 6**. Frequency histograms reflecting distributions of raw trials of latencies (ms) available for each site (I-VII) across all neonates in the ASD and the non-ASD cohorts.
- **Supplementary Figure 7.** Pairwise comparison of site x group to measure the similarity between distributions estimated with the bootstrapping technique (equal number of points in each set).
- **Supplementary Figure 8.** Personalized approach localizing each neonate on a parameter space spanned by the features and statistical parameters empirically estimated from the micromovements data type.
- **Supplementary Figure 9.** Fluctuations in peaks' prominences separate dB levels in ASD and non-ASD pre-term and full-term neonates, and automatically distinguish for each level, the different groups.
- **Supplementary Figure 10.** Empirically estimated continuous Gamma family of probability distribution parameters (shape and scale) of minimum and maximum peak amplitude values (μV) in response to three levels of clicks ranging from 70-80dB.
- **Supplementary Figure 11.** Empirically estimated continuous Gamma family of probability distribution parameters (shape and scale) of minimum and maximum peak prominence (μV) values in response to three levels of clicks ranging from 70-80dB.
- **Supplementary Figure 12.** Empirically estimated continuous Gamma family of probability distribution parameters (shape and scale) of minimum and maximum peak widths (ms) values in response to three levels of clicks ranging from 70-80dB
- **Supplementary Figure 13.** Pairwise comparison of cohorts upon bootstrapping to obtain equal number of trials per group and taking the median outcome value across comparisons.
- **Supplementary Figure 14.** Distributions of neonates across the Gamma moments parameter space for the three features under consideration and for trials drawn from each of the responses to 70-75-80dB clicks' level.

- **Supplementary Figure 15.** Parameter space spanned by the empirically estimated Gamma parameters and EGA.

### Information about participants

**Neonates' Auditory Brainstem Response (ABR)—** The first set involving 233,917 newborn hearing screening records from NICU *vs.* WBN uses a sampling rate of 5kHz (0.2 ms period) which produces a compressed ABR waveform of 62 data points for the right ear and 61 for the left (12.4ms window) - the right and left ears are stimulated simultaneously at slightly different rates to extract the ear specific data. These data do not contain any pre-stimulus data. Midline vertex-to-ape recordings were performed using 100 $\mu$sec rectangular rarefaction click stimuli 35 dB above adult normal hearing level (nHL). Stimulus delivery and response averaging were done using Intelligent Hearing System's ABR-based SmartScreenerPlus hearing screening system. The peak V latency used in the analysis for the right and left ears respectively is also provided, thus we can obtain the interaural differences.

The second set of neonates had ipsilateral left ear vertex-to-mastoid recordings performed using 100 $\mu$sec rectangular rarefaction click stimuli 80 dB nHL presented at a rate of 12.9/s. Our standard average ABR waveform consisted of 3,072 (3 sets of 1,024) artifact-free trials. Two additional sets of 2,048 (2 sets of 1,024) trials were recorded at 75 and 70 dB nHL. Stimulus delivery and response averaging were done using Intelligent Hearing System's ABR SmartEP system. Multiple full L1 sets comprising all trials of three different dB levels were available for all babies and additional trials L2-L3 were available for some babies. These are reflected in Table 1. These data set from ABRs (at least 3 trials each) were administered during the newborn period. For NICU infants they typically occurred at the bedside shortly after birth, but no earlier than 24 hours postnatally and 31–32 weeks post-conceptional age. (Mean (SD) age at test was 37.5 (4.1) weeks). This cohort comprised 64.9% males/35.6% females and were ethnically diverse. They were 31.2% Latinx (including Latinx Black, White, Asian, and multiracial) and 68.8% non-Latinx, with 16.9% Black; 54.5% White; 3.9% Asian, 1.3% Indian, and 23.4% multiracial. Data were collected at the Richmond University Medical Center, US (NICU babies) and at New York State Institute for Basic Research in Developmental Disabilities (WBN). The same personnel collected the data with identical instrumentation and settings.

These participants are part of our longitudinal study of attention and arousal in children at-risk for developmental problems due to pre-term birth and/or medical factors requiring assignment to the NICU (*22*). Selection criteria for the larger study included any of the following: birthweight (BW) (<1,800 g); fetal distress with evidence of asphyxia at birth; assisted ventilation (>48 h); persistent apnea or bradycardia; abnormal neurological signs; small for gestational age (<10th percentile BW for gestational age), intrauterine growth restriction, or dysmature; multiple gestation (at least one twin met criteria or BW <2,000 g). Exclusion criteria were known prenatal exposure to drugs of abuse, diagnosis of a major congenital anomaly, or chromosomal disorders. In addition, a sample of healthy newborns, recruited from the well-baby newborn nursery (WBN), participated in the original study as controls. Additional criteria for the current study were: (1) Administered an ABR during the newborn period; (2) Behaviorally assessed at older ages to ascertain or rule out an ASD diagnosis, (3) Absence of significant brain damage documented by cranial ultrasound, CT, MRI such as IVH (Papile Grade >I), ventriculomegaly (>5 mm), periventricular leukomalacia, hydrocephalus, or seizures requiring treatment documented by EEG.

The data analyzed pertained to a subset of children from this study, with *n=24* receiving an ASD diagnosis (18 NICU; 6 WBN). Diagnoses were determined as follows: by clinicians at the Institute for Basic Research in Developmental Disabilities (*n=9*); through participation in Early Intervention or special preschool programs (*n=8*); by parent interview with the ADI-R (*n=6*); by a private neurologist (*N=1*). A control group comprised of 30 children was selected (19 NICU; 11 WBN). No child in the control group was diagnosed with ASD or was suspected of being on the autism spectrum, based upon behavioral testing at their 24/36-month follow-up assessments. In addition, based upon the PDD Behavior Inventory (*23*), none were reported to show any signs of ASD by their mothers; none were receiving special services because of ASD or suspected ASD.

| **ABR TEST 1 (L1)** | | | | |
|---|---|---|---|---|
| | **NICU PT** # babies | **NICU FT** # babies | **WBN (FT)** # babies | **TOTAL** # babies |
| **ASD** | 16 | 2 | 6 | 24 |
| **Non ASD** | 16 | 3 | 11 | 30 |
| **TOTAL** | 32 | 5 | 17 | 54 |
| **Additional Trials ABR TEST 2 (L2)** | | | | |
| **ASD** | 3 | 1 | | 4 |

| | | | | |
|---|---|---|---|---|
| **Non ASD** | 3 | 1 | | 4 |
| **TOTAL** | 6 | 2 | | 8 |
| **Additional Trials ABR TEST 3 (L3)** | | | | |
| **ASD** | 4 | | | 4 |
| **Non ASD** | | | | |
| **TOTAL** | 4 | | | 4 |

**Supplementary Table 1**. Number of babies included in the study and sets L1, L2, L3 of repeated trials. Each trial contained three levels of sound spanning 70, 75, 80dB from which the full waveform was saved.

**Young Infants and Children's Auditory Brainstem Response (ABR)—**In the third set of young infants and young children, multiple-trial ABR recordings per participant were acquired from 65 individuals, 18 with an ASD diagnosis and 47 typically developing controls. In the ASD, we have 13 males and 5 females. In the neurotypicals, we had 23 males and 24 females. These data were obtained with a sampling resolution of 40kHz (spaced 25µs) using stimulus of 65 dB nHL at a stimulation rate of 19.3/second. Male and female participants ranged from 1.8 to 6.8 years if age. There were multiple trials per infant and child, totaling 215 trials.

**Good Quality Data Available for Empirical Estimation Distributional Analyses**

We had access to 108 trials from 54 babies and L1, L2, L3 full sets containing the three dB levels (70, 75, 80 dB). Among these, 66 had some repeated trials, (full set of 3 dB levels each), 54 L1; 8 L2 and 4 L3. These data are detailed in Table 1 for each baby type.

We had 54 babies with sufficient data (over 100 peaks per combined waveforms) for personalized analyses that included empirical distributional parameter estimation (with 95% confidence interval criterion) and a unique trial L1 encompassing all 3 dB levels. For population analyses, we pooled across all three trials (L1, L2, L3) addressing distinctions in ABR amplitudes and latencies for each baby type.

We also had access to additional trials with full ABR waveform data from 14 FT babies (2 FT NICU babies) and 34 NICU PT babies. However, these data sets were incomplete (*e.g.,* missing data from a dB level or two) or too noisy. As such, these data were not included in the present analyses.

In addition to the full waveform data ($\mu V$), we also had access to the latency data (*ms*) from 47 unique recording sessions including the following neonates (13 FT, 11 FT NICU, 2 FT nonNICU with 8 ASD and 34 PT with 30 ASD). As with the full waveform, we had multiple trials in each set of responses to three dB levels. Pooling across all neonates and trials, and using bootstrapping techniques, we built a larger dataset (above 100 measurements) to interrogate fluctuations in latencies and various features of the full waveforms.

### *Clinical Data*

Infants were recruited in the span of 17 years, some prior to 2011 and some after 2011. Below we detail the clinical tests.

**Clinical Tests:** Beginning at 3 months of age, and extending through 25 months, infants participating in the study prior to 2011, were tested with the Bayley Scales of Infant Development - 2nd Edition (BSID-II) (Bayley, 1993) measuring cognition, motor, and behavioral development. It has been used widely both in research and clinical applications and is currently in its 4th edition. It comprises Mental, Motor, and Behavior Rating Scales. Only the first two scales were analyzed in the current study. Age-appropriate norms were applied to the scaled scores to derive a Mental Developmental Index (MDI) and a Psychomotor Developmental Index (PDI). Both indices have a mean of 100 and a standard deviation of 16.

Infants participating in the study after 2011 were assessed from 3- through 24-months of age with the Mullen Scales of Early Learning (MSEL AGS Edition, 1995). The MSEL is a comprehensive measure of cognitive and motor functioning divided into five areas including Gross Motor, Visual Reception, Fine Motor, Receptive and Expressive Language. It yields separate scores in each of these domains as well as a composite Standard Score made up 4 of the 5 areas (Gross Motor is excluded).

### Population Analyses from Pooled Data and Bootstrapping

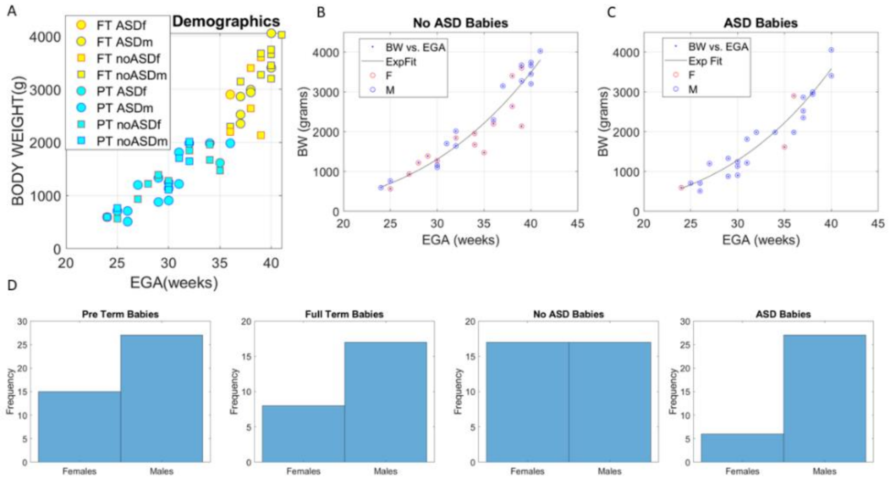


**Supplementary Figure 1. Sample cross-sectional demographic graphs of neonates reveals nonlinear relations between early weight growth and estimated gestational age**. Body weight expressed as a function of estimated gestational age (EGA), color coded as full-term (yellow FT) or pre-term (cyan PT) categories. Marker edge denotes sex (red female, blue male) and marker type denotes diagnosis type (circle ASD, square non-ASD). (B-C) Nonlinear relation between estimated gestational age (EGA) and body weight (BW). (D) Available *trials* from neonates with complete demographic records available per baby type used in supplementary material figures featuring population statistics.

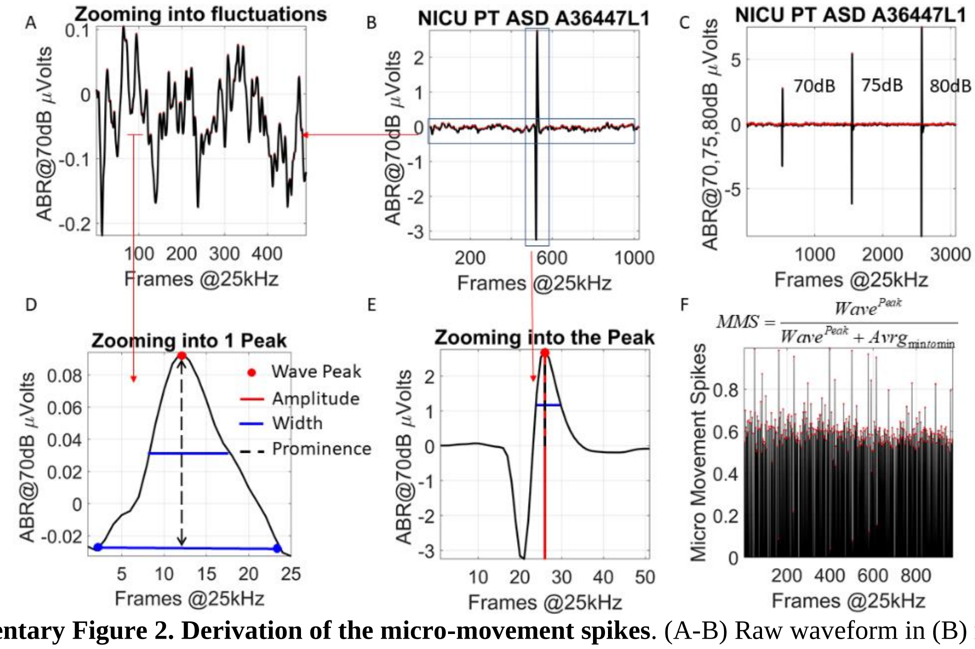


**Supplementary Figure 2. Derivation of the micro-movement spikes**. (A-B) Raw waveform in (B) represents the auditory brainstem response ABR from one click at *e.g.,* 70dB. (C) Sample ABRs from three clicks at 70, 75 and 80dB. (D) Each pulse (Dirac delta) is preceded and followed by microscopic jitter that are zoomed in panel (A). The full waveform is thus obtained, and the amplitudes (μV) and latency of the peaks (jitter and Dirac delta combined) used to scale as in panel D, where we show the peak amplitude, width, and prominence features under consideration. These features are extracted from both the jitter and pulse. (E) Zooming into the pulse to highlight the features under consideration in (D). (F) Resulting micro-movement spikes derived from the jitter and pulse features (in this case for the prominences) using the formula in the panel to scale out allometric effects due to anatomical differences that emerge from different body sizes and masses rapidly changing in early infancy. The micro-movements data type can be derived from any time series by obtaining the absolute deviations from the mean amplitude or inter-peak-interval timings, or any other feature of the waveform that takes then the series of fluctuations away from the mean and scales them appropriately between 0-1. This standardized waveform can then be used in further analytical pipelines to compare participants of disparate ages and anatomical types.

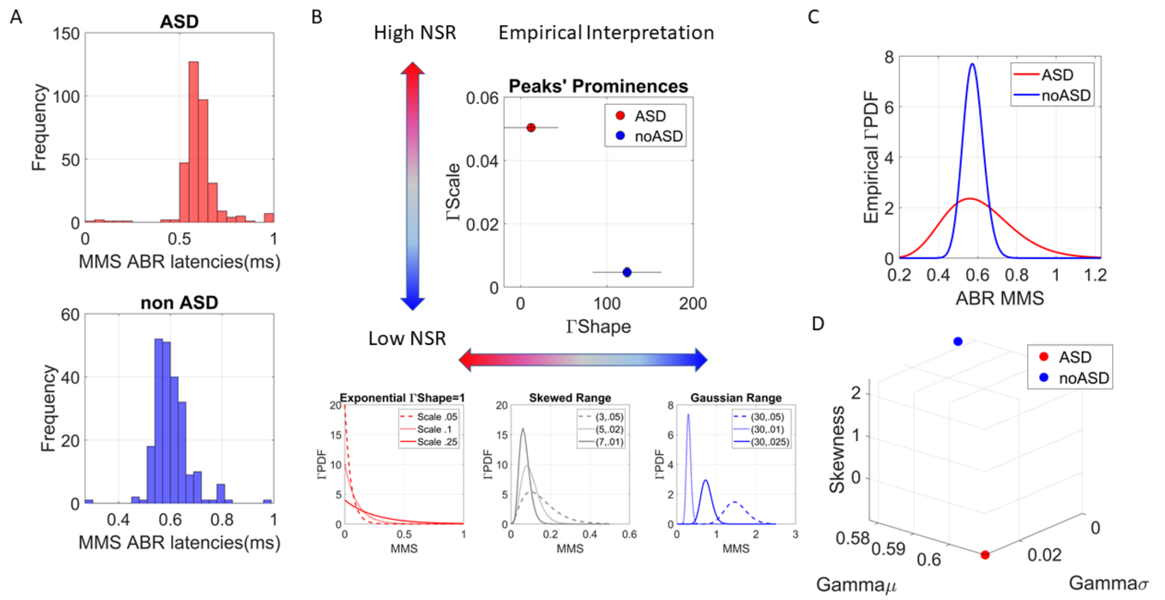


**Supplementary Figure 3. Empirical estimation and inference of stochastic signatures derived from the micro-movement spikes**. (A) Sample frequency histograms of micro-movement spikes derived from the fluctuations in the peaks' latencies (*ms*) normalized to be unitless as in Figure 12. (B) The Gamma parameter plane spanned by the shape and scale parameters empirically estimated using maximum likelihood estimation (MLE). The estimated parameters are plot with 95% confidence intervals for each estimated shape and scale value. These two data points correspond to the PDFs in (C). Gradient colored arrows denote the empirical ranges of human micro-movements PDFs, as previously obtained from different biosensors across different systems and levels of control, in several thousand human participants across the lifespan. Notice that because the MMS are standardized and scale out anatomical disparities, we can place points from different participants with different anatomical features (due to different ages) on the same parameter space. Along the shape dimension, values range from 1 (Exponential distribution representing a memoryless random process and being a special case of the continuous Gamma family) to over 100, for the Gaussian distribution-like symmetric case, with skewed distributions falling in between these two extreme cases. Along the scale (dispersion) dimension, lower values represent lower noise to signal ratio (NSR). This is so because given the Gamma variance and mean, and given the Gamma shape ($a$) and the Gamma scale ($b$), $NSR = \frac{a \cdot b^2}{a \cdot b} = b$ (D) The empirically estimated Gamma moments can be obtained as well and represented on a parameter space spanned by the mean (x-axis), the variance (y-axis), the skewness (z-axis) and the kurtosis proportional to the size of the marker. Lower NSR (blue non-ASD) has lower variance in this case. Given the empirically estimated MMS values, we can make well informed inferences with highly interpretable value about the stochastic signatures. This is so because we have characterized them in humans before.

| Regions | I | II | III | IV | V | VI | VII | Cum Σ |
|---|---|---|---|---|---|---|---|---|
| ASD | 1.68 | 3.58 | 4.80 | 6.17 | 7.29 | 8.48 | 9.68 | |
| Δ ASD | | 1.90 | 1.22 | 1.37 | 1.12 | 1.19 | 1.20 | 8.00 |
| Non-ASD | 1.66 | 3.33 | 4.78 | 5.83 | 6.95 | 8.11 | 9.28 | |
| Δ Non-ASD | | 1.67 | 1.45 | 1.05 | 1.12 | 1.16 | 1.17 | 7.62 |
| Interpeak Δ (Non-ASD - ASD) | 0.02 | 0.25 | 0.02 | 0.34 | 0.34 | 0.37 | 0.40 | 1.74 |

Supplementary Table 2. Peak latencies for each site in the two cohorts and inter-peak latency differences. Last column shows the cumulative sum of the delays and the appreciable ASD neonates' delay of 1.74 *ms* (considering that sound processing is generally agreed to be on a $\mu s$ time scale)

### Supplementary Figure 4

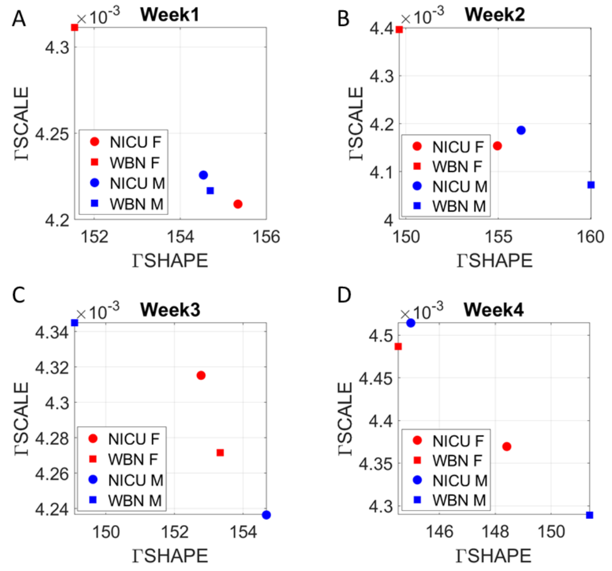


**Supplementary Figure 4. Tracking neonates from NICU and WBN cross-sectionally 4 weeks reveals differentiation between males and females withing and between each group.** Stochastic signatures of empirically estimated Gamma distribution shape vs. scale (signaling dispersion and noise to signal ratio) reflect the variations in the minimum latency of Peak V across a random draw of 100 babies in each week and group (NICU Female, NICU male, WBN Female and WBN Male.)

### Supplementary Figure 5

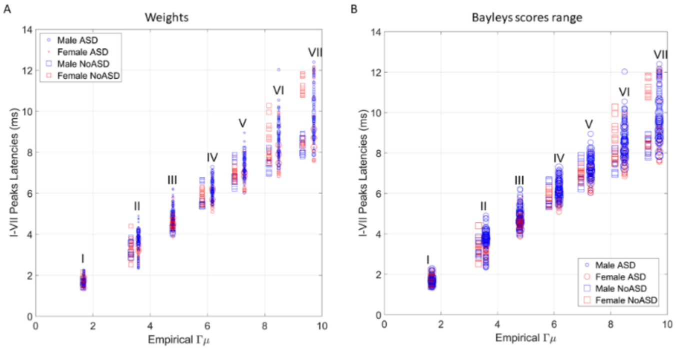


**Supplementary Figure 5. Empirically estimated Gamma means (x-axis) *vs.* all latencies (*ms*) for each site (I-VII) show the spread of values across babies in each group**. Marker size is proportional to clinical measure value, marker shape denotes ASD (circle) or non-ASD (square), and color denotes sex in legend (A) Body weight at the first visit (smaller size is lower weight) ranging from 600 to 4,000 grams (see Figure 10 in the methods of the main paper). (B) Bayleys scores proportional to the size of the marker.

### Supplementary Figure 6

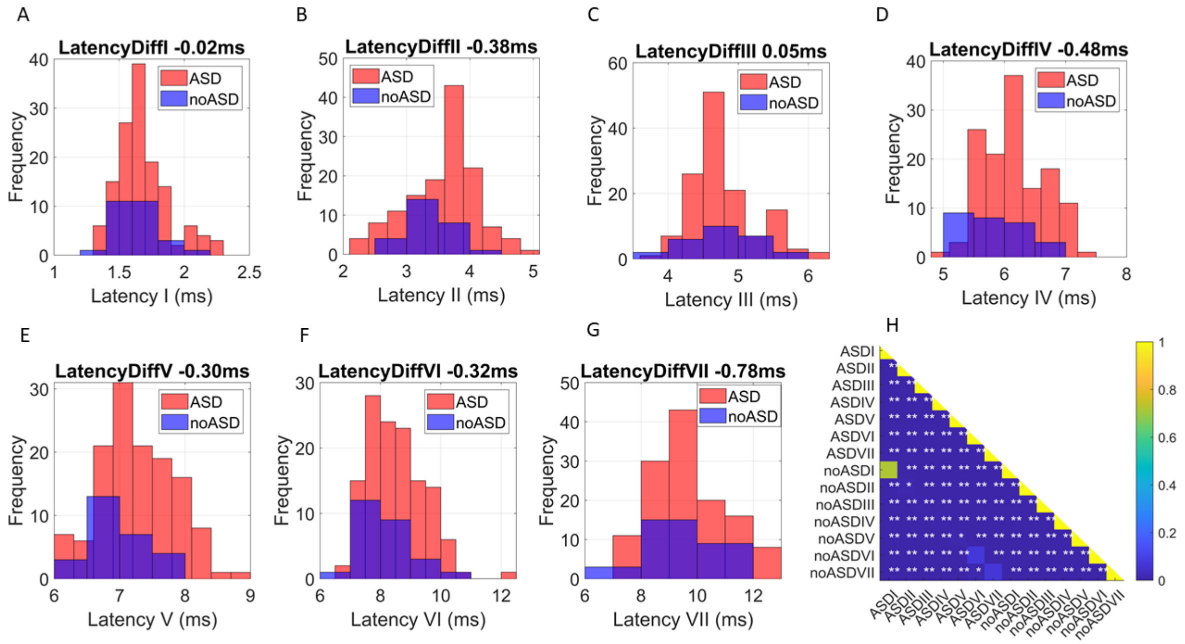


**Supplementary Figure 6 Frequency histograms reflecting distributions of raw trials of latencies (*ms*) available for each site (I-VII) across all neonates in the ASD and the non-ASD cohorts.** (A) Site I at the ear has a shift of 0.02ms delayed peak in the ASD cohort. (B-D) Sites II, III and IV at the pons shift 0.38, 0.05, 0.48ms respectively, with delayed peaks in the ASD cohort. (E-F) Sites V and VI of the midbrain have 0.30ms and 0.32ms delays respectively, for the ASD cohort. (G) Site VII at the primary auditory cortex has a 0.78ms delay of the peak latency in the ASD cohort. (H) Bootstrapping by drawing from the larger set the number of measurements of the smaller set and forming 100 distributions compared with non-parametric tests and obtain the median p-value to construct a pairwise matrix of comparisons for each site (7 sites) and group (2 groups). This matrix is color-coded and entries with two asterisks are significant $p < 0.01$, while those with one asterisk are $p < 0.05$.

**Supplementary Figure 7**

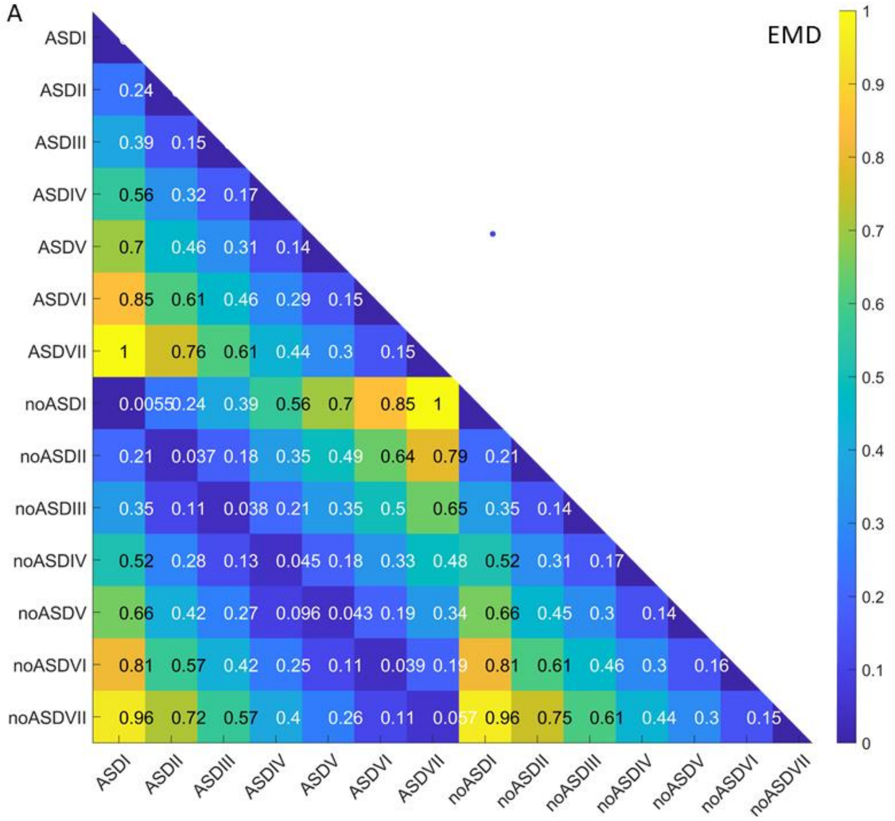


**Supplementary Figure 7, Pairwise comparison of site x group to measure the similarity between distributions estimated with the bootstrapping technique (equal number of points in each set).** We use the Earth Mover's Distance to ascertain the similarity between distributions and normalize the values to color the matrix entry accordingly. Notice the cumulative nature of the delays in the color gradient thus obtained.

### Personalized Analyses

Supplementary Figure 5 represents a parameter space spanned by the skewness of the distribution of peaks' width along the *x*-axis, the variance of the distribution of peak's amplitude along the *y*-axis and included the body weight at the first visit (panel A), or the estimated gestational age (EGA) at birth (panel B) along the *z*-axis. Each point in this parameter space represents one baby (personalized stochastic signatures) along each estimated family of probability distributions spanned by the waveforms' features. The graph shows automatically emerging clusters of babies. This clustering reveals that low birth weight (BW) and low estimated gestation age, EGA are not predictors of ASD. The cohort of ASD neonates had babies across low and high values of both parameters.

**Supplementary Figure 8**

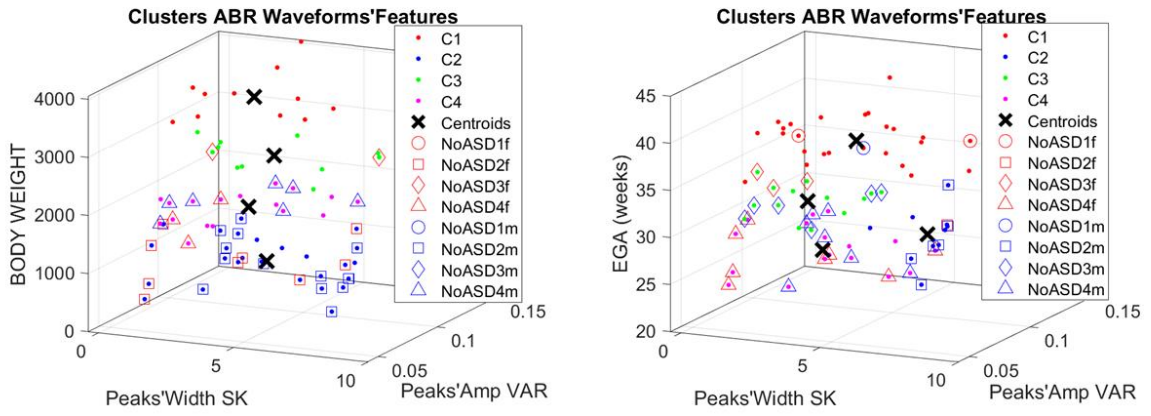


**Supplementary Figure 8. Parameter space spanned by the features and statistical parameters empirically estimated from the micromovements data type.** The peaks' width is along the x-axis, the peaks' amplitude runs along the y-axis and the body weight runs along the z-axis (left panel) or the estimated gestational age (right panel). K-means with 4 clusters groups non-ASD on lower ranges of weight, while ASD spans all weight values. Likewise, EGA spans in ASD babies all values. Legend shows the symbols and color denoting sex and diagnosis for each cluster. These results reveal that weight and EGA are not predictors of ASD during neonatal stages of development.

### Waveform's prominences separate ASD from non-ASD in full-term vs. pre-term babies

The waveforms reflecting the brainstem responses had never been analyzed in their full extent, as only the peaks' averages had been of interest to prior research (*27*, *28*). In addition to the group analyses, we used a personalized approach that empirically estimates the stochastic signatures of ABR parameters for each member of the cohort. We did so in search of self-emerging patterns along parameter spaces amenable to automatically stratify the cohort and identify statistical features maximally differentiating each clinical group. ABR latencies at *ms* time scale, along with the fluctuations in the waveforms' peaks' amplitude ($\mu V$) and peaks' width (*ms*) reflected marked, unambiguously different responses between the ASD and non-ASD subgroups.

The individual differences are shown in Supplementary Figure 5, while Supplementary Figure 6 shows the group results. There we see different stochastic signatures in response to each of the dB levels under consideration. Not only did the ASD deviate from the non-ASD in the peaks' latencies, but they also depart in their fluctuations in peaks' amplitudes, widths, and prominences. For each of these features, the PDFs are different. And they are different for each dB level.

Although these differences extend through all parameters under consideration, we identified the peaks' prominences as the feature revealing the largest differentiation in PDFs between these two cohorts, *i.e.*, maximally separating ASD from non-ASD full-term neonates across the dB levels.

This waveform's feature also systematically differentiated ASD pre-terms from non-ASD pre-terms (Supplementary Figure 6A shows the separation on the log-log Gamma parameter plane, while B does so for the empirical PDFs. Panel C shows the corresponding empirical Gamma moments along a path connecting the signatures for each dB level's response. We can automatically separate these cohorts along all three dB levels under consideration and distinguish pre-term from full-term neonates according to the ASD *vs*. non-ASD subtype. All other pairwise non-parametric comparisons (Wilcoxon rank sum test) yielded statistically significant differences for these three dB levels' differentiation at the *0.05* alpha level.

### Supplementary Figure 9

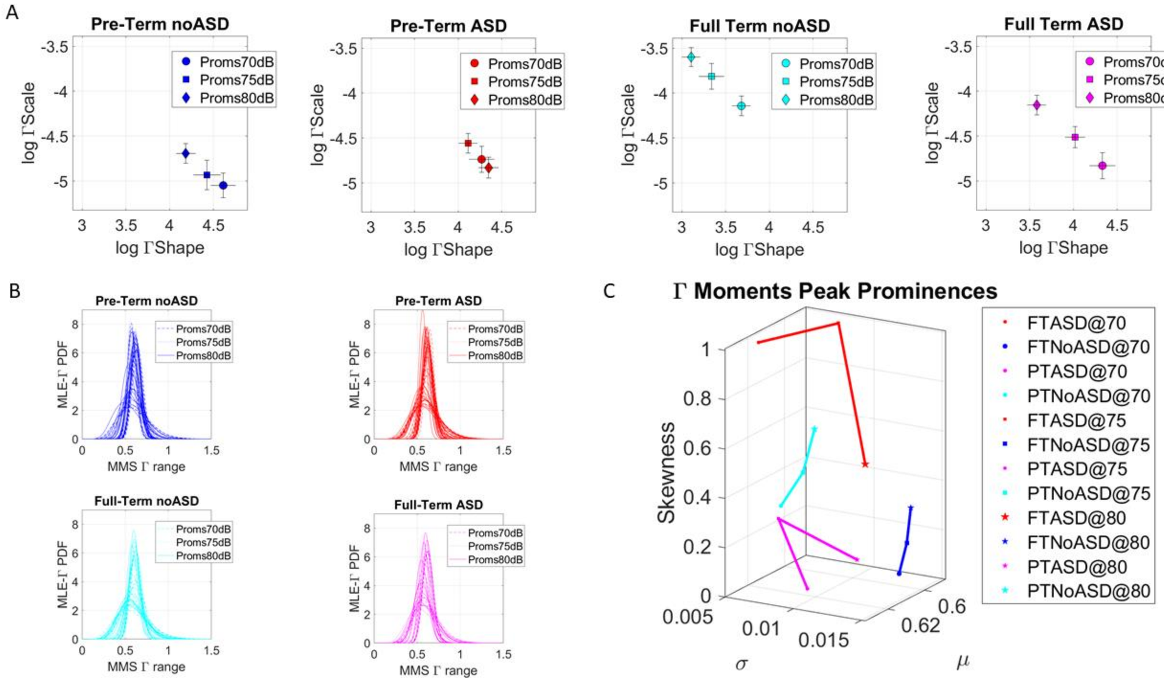


**Supplementary Figure 9. Fluctuations in peaks' prominences separate dB levels in ASD and non-ASD pre-term and full-term neonates, and automatically distinguish for each level, the different groups.** (A) Log-log Gamma parameter plane contains the stochastic signatures for each dB level (with 95% confidence intervals). These levels are different for each of the ASD *vs*. non-ASD cases and pre-term *vs*. full-term cases, with the later showing maximal separability between ASD and non-ASD neonates. (B) Corresponding empirical Gamma PDFs obtained for each baby that went into the computation of the Gamma shape and scale group parameter. (C) Empirical Gamma moments spanning parameter space along the x-axis (mean), y-axis (variance), z-axis (skewness) and using the kurtosis proportional to the marker's size. The stochastic signature of each dB level connects to the next level via a line representing a path in probability space. This path clearly marks differentiation in the transitions of the signal from one dB level to the next in ASD *vs*. non-ASD neonates.

Supplementary Figures 10-12 show results for the peaks' amplitude, prominences, and widths, while Supplementary Figure 13A captures differences in frequency histograms obtained through the earth mover's distance metric (EMD) matrices. This matrix has entries denoting pairwise similarity in amplitude and inter-peak-interval latencies of the full waveform, while Supplementary Figure 13B shows the results from non-parametric pairwise comparisons of the differences in peak amplitude values for each subgroup of the 70-75-80dB -prominences (left) and amplitudes (right). There the differentiation of three dB levels in full-term *vs*. pre-term ASD neonates are statistically indistinguishable. This means that the fluctuations in the differences in peaks' prominences across the levels cannot separate between pre-term and full-term ASD babies.

### Supplementary Figure 10

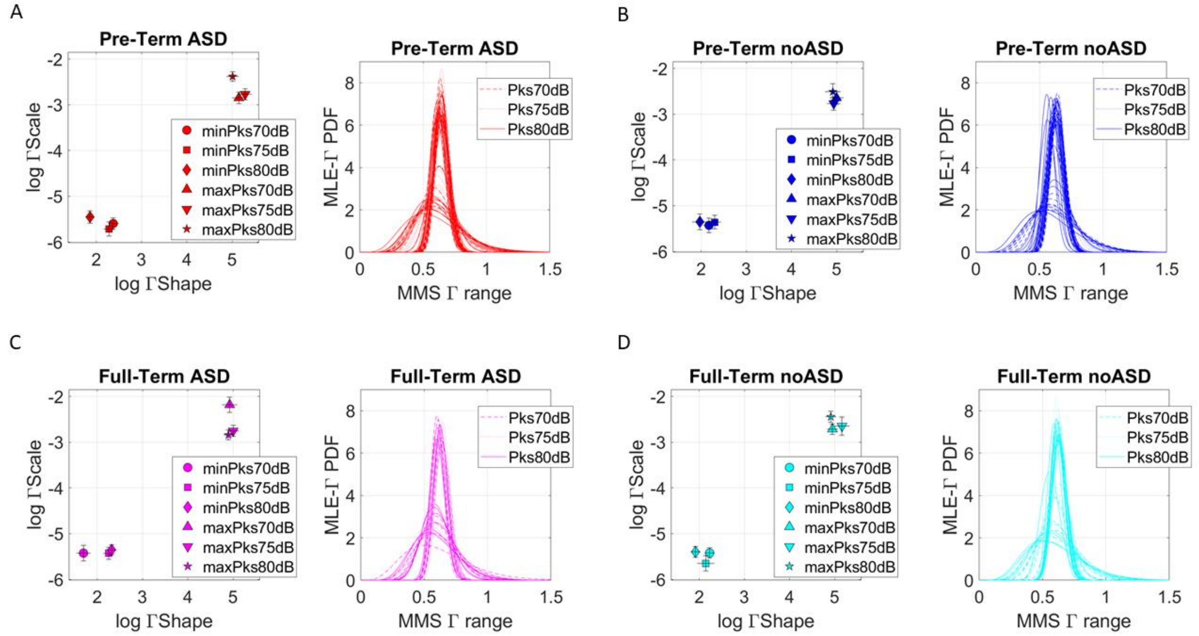


**Supplementary Figure 10. Empirically estimated continuous Gamma family of probability distribution parameters (shape and scale) of minimum and maximum peak amplitude values (μV) in response to three levels of clicks ranging from 70-80dB.** Plots show the range of PDFs across newborns, ASD *vs.* non-ASD and full-term (A-B) *vs.* pre-term (C-D).

### Supplementary Figure 11

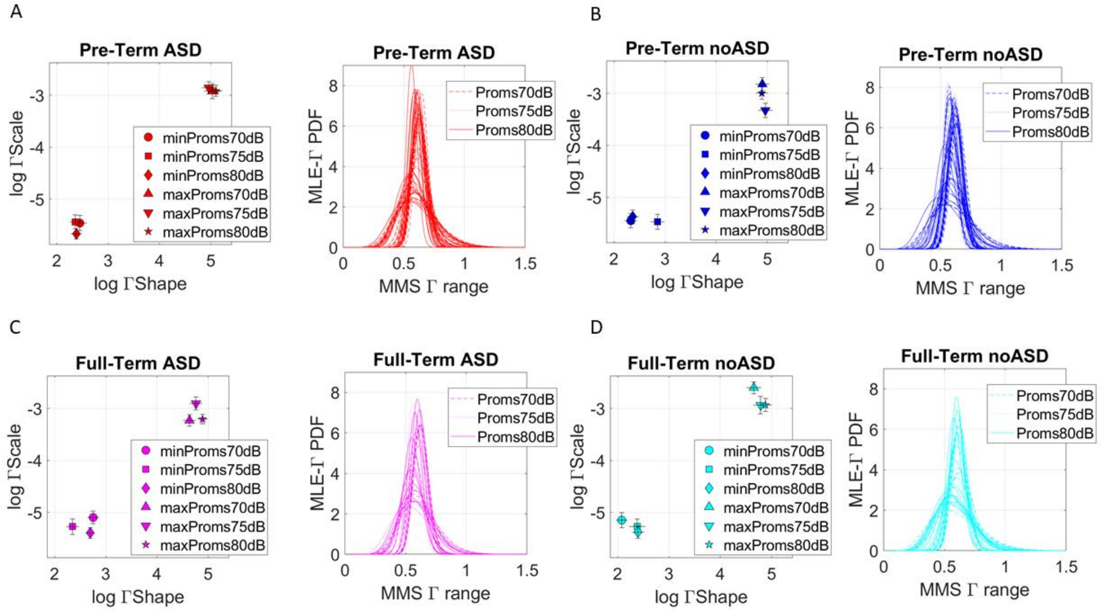


**Supplementary Figure 11. Empirically estimated continuous Gamma family of probability distribution parameters (shape and scale) of minimum and maximum peak prominence (μV) values in response to three levels of clicks ranging from 70-80dB**. Plots show the range of PDFs across newborns, ASD *vs.* non-ASD and full-term (A-B) *vs.* pre-term (C-D).

### Supplementary Figure 12

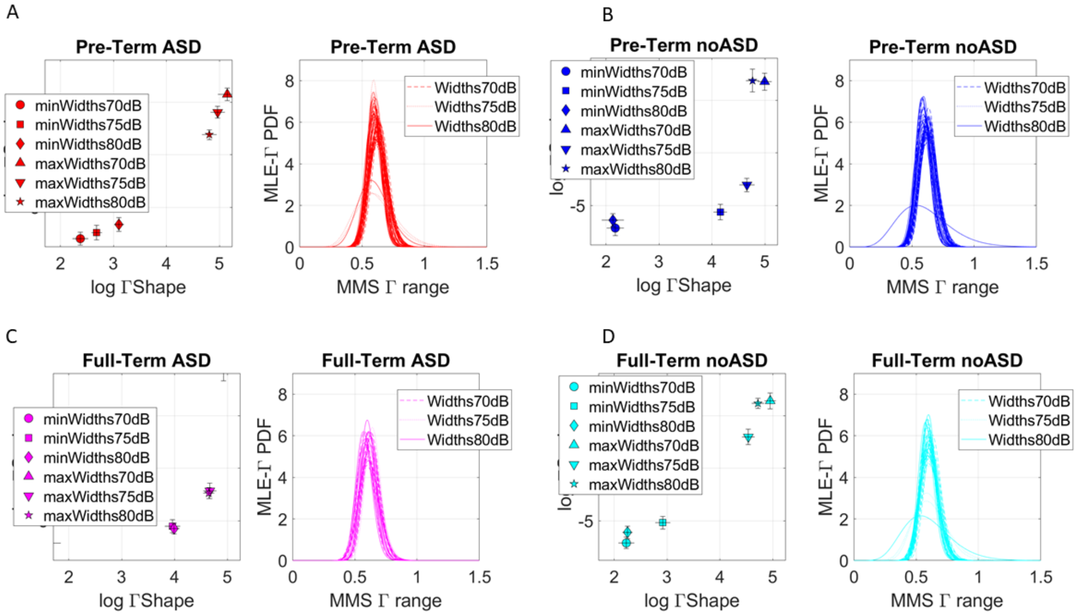


**Supplementary Figure 12. Empirically estimated continuous Gamma family of probability distribution parameters (shape and scale) of minimum and maximum peak widths (*ms*) values in response to three levels of clicks ranging from 70-80dB**. Plots show the range of PDFs across newborns, ASD *vs.* non-ASD and full-term (A-B) *vs.* pre-term (C-D).

### Supplementary Figure 13

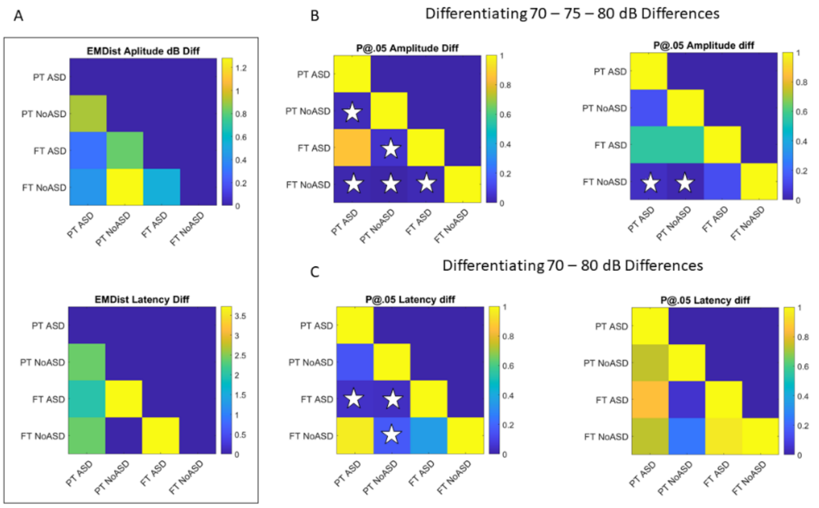


**Supplementary Figure 13. Pairwise comparison of cohorts upon bootstrapping to obtain equal number of trials per group and taking the median outcome value across comparisons.** (A) EMD similarity metric color coded to show differentiation across groups (lower values blue show more similarity) for amplitude (top) and latency or inter-peak-interval timings (bottom). (B) Difference values between 70-75-80 dB are pairwise compared to ascertain statistically significant differences pairwise. (Left panel) Fluctuations in peaks' prominences differences across click levels are indistinguishable across ASD neonates regardless of whether they are pre-term or full-term, but all other comparisons are significant at the 0.05 alpha level according to the non-parametric Wilcoxon rank sum test. (Right panel) Fluctuations in peaks' amplitudes are significant at the alpha 0.05 level for full-term non-ASD and all pre-term (ASD and non-ASD) neonates. (C) Comparison of 70-80dB differences using the latencies (ms) of the inter-peak intervals of the prominence series (left panel) and the inter-peak-interval values of the full amplitude series.

### Supplementary Figure 14

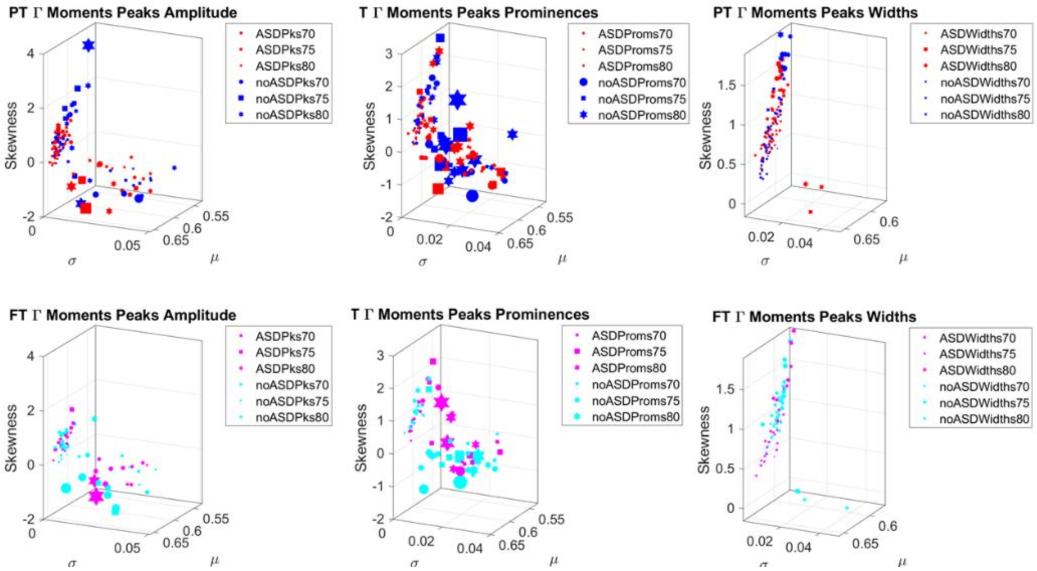


**Supplementary Figure 14. Distributions of neonates across the Gamma moments parameter space for the three features under consideration and for trials drawn from each of the responses to 70-75-80dB clicks' level.** Legend shows the diagnosis type and level for each feature of the waveform.

### Supplementary Figure 15

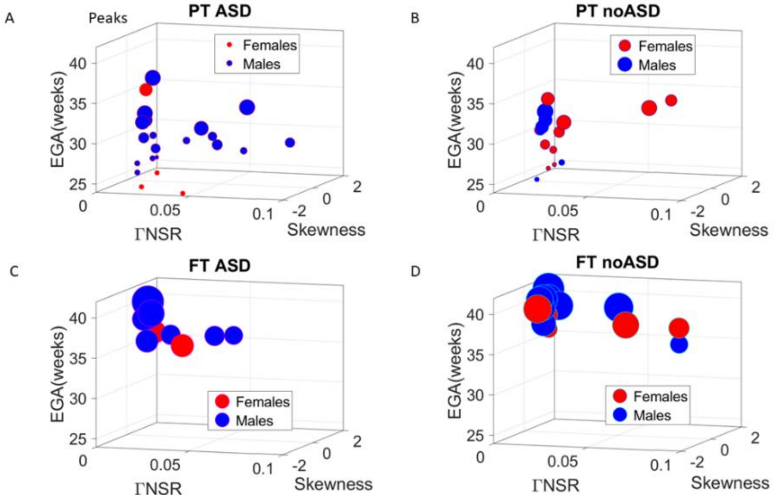


**Supplementary Figure 15. Parameter space spanned by the empirically estimated Gamma parameters and EGA** (x-axis scale (noise to signal ratio), y-axis skewness and z-axis EGA for the peaks' amplitudes (μV) and for each subgroup of pre-term *vs*. full-term babies, and ASD *vs*. non-ASD babies. The size of the marker is proportional to the weight (spanning from 600-4000 grams).
